# The GNMT N-terminus Couples Folate Feedback to Methyl-donor Homeostasis

**DOI:** 10.64898/2026.05.26.727678

**Authors:** Isaac Kraz, Oi Wei Mak, Subray Hegde, Benjamin M. Lorton, Harrison Hector, Michael K. Broussalian, Sarah Graff, Jennifer Aguilan, Jeffrey Bonanno, Roozbeh Eskandari, Daniel Groom, Simone Sidoli, Steve Almo, Michael Brenowitz, Evripidis Gavathiotis, Derek M. Huffman, David Shechter

**Affiliations:** Department of Biochemistry, Albert Einstein College of Medicine, Bronx, NY 10461; Department of Molecular Pharmacology, Albert Einstein College of Medicine, Bronx, NY 10461; Institute for Geroscience, Albert Einstein College of Medicine, Bronx, NY 10461; Department of Medicine, Albert Einstein College of Medicine, Bronx, NY 10461

**Keywords:** One Carbon Metabolism, GNMT, Methionine, Folate, SAM, Methylation

## Abstract

Maintenance of S-adenosylmethionine (SAM) homeostasis is essential for methylation of biomolecules, nucleotide and polyamine synthesis, and redox homeostasis. While all methyltransferases consume SAM, only a subset of highly tissue specific methyltransferases regulates SAM homeostasis. Among them, glycine N-methyltransferase (GNMT) is enriched in the liver and its dysregulated activity has been linked to compromised liver function. GNMT is inhibited by the methyl carrier 5-methyltetrahydrofolate (5mTHF), suggesting a negative-feedback mechanism regulating its activity. Here, we identify the GNMT N-terminal tail, and specifically phosphorylation at serine 9 (S9ph), as a regulatory modification linking folate-dependent feedback inhibition to SAM homeostasis. Structural and biochemical analyses and molecular dynamics simulations revealed that the N-terminal tail is required for catalytic turnover of SAM and for 5mTHF binding. Phosphoproteomic analysis showed that GNMT S9ph is abundant in mouse liver and further enriched in aged mice. Consistent with loss of folate-dependent negative feedback, both distal N-terminal truncation (residues 1-8) and a phosphomimetic substitution abolished 5mTHF binding while maintaining catalytic activity. In hepatocyte cell lines lacking endogenous GNMT, lentiviral overexpression of constitutively active GNMT mutants depleted SAM, increased SAH, disrupted protein methylation, impaired growth, and induced transcriptional responses consistent with methyl-donor stress. Together, these findings identify the GNMT N-terminus as a tunable phosphoregulatory domain that dynamically regulates GNMT activity and cellular methylation potential.

## Introduction

One carbon metabolism, composed primarily of the methionine and folate cycles, is a critical and highly regulated pathway involved in the production and transfer of one carbon units involved in methylation reactions, purine synthesis, and glutathione production. S-adenosylmethionine (SAM), a product of one carbon metabolism, serves as the universal methyl donor for methyltransferase enzymes, which catalyze transfer of the sulfonium methyl group of SAM to nucleophilic acceptors such as the ε-amino group of lysine residues on proteins^1–3^, the 5-position of cytosine in DNA^4–6^ and RNA^7,8^, or the ethanolamine head group of phosphatidylethanolamine in lipids^9,10^. These reactions generate S-adenosylhomocysteine (SAH), which is converted to homocysteine by the enzyme S-adenosylhomocysteine hydrolase (AHCY). Homocysteine can either enter the trans-sulfuration pathway which generates antioxidant species such as hydrogen sulfide and glutathione or may be remethylated by methionine synthase to regenerate methionine.

Precise regulation of cellular SAM pools is critical to maintain cellular methylation potential (often measured by the SAM:SAH ratio), where disruption of this potential leads to broad cellular perturbations including altered DNA^11–13^ and histone methylation^14,15^, transcriptional dysregulation^14,16,17^, and impaired cell growth^18–21^. In the disease context, reprogramming of one carbon metabolic pathways and disruption of SAM:SAH homeostasis contributes to multiple disease states, including hepatic steatosis often precipitating neoplastic changes^22, 23,24^ and aging^25–31,23^. Cancer cells are especially dependent on one-carbon metabolism to support cell division, since purine and pyrimidine biosynthesis branch from this pathway. Folate-pathway genes such as TYMS and DHFR are upregulated in cancer and form the basis for antifolate chemotherapeutic strategies^22,33^.

While all methyltransferases consume SAM, only a few substantially alter cellular SAM pools. Among these, nicotinamide N-methyltransferase (NNMT)^33^, phosphatidylethanolamine methyltransferase (PEMT), and glycine N-methyltransferase (GNMT) have previously been shown to act on macromolecular and metabolic substrates. Glycine N-methyltransferase (GNMT) catalyzes the SAM-dependent methylation of glycine, forming sarcosine. Prior studies show that GNMT modulates SAM abundance, where GNMT knockout mice had elevated SAM and SAM:SAH ratios and, conversely, overexpression of GNMT in cancer cell lines led to a depletion of SAM. GNMT is inhibited by 5-methyl-tetrahydrofolate (5mTHF), an abundant folate pathway metabolite^34^. 5mTHF-mediated inhibition has been demonstrated *in vitro* to involve the GNMT N-terminus, a highly conserved domain composed of a flexible tail (residues 1-8) followed by a structured U-loop (residues 9-22) proximal to the globular domain and SAM binding site, the latter of which functions as a gatekeeper for SAM entry^35,36^.

GNMT perturbation affects tissue- and organism-level homeostasis, however the mechanisms underlying these changes are poorly understood. Decreased GNMT activity and expression are associated with liver specific pathologies including NAFLD/NASH and hepatocellular carcinoma^39–44^. These observations demonstrate the importance of GNMT regulation and support our efforts to better understand these mechanisms.

Here, we identify the GNMT N-terminus as a dynamic regulatory domain linking folate sensing to methyl-donor homeostasis. Using biochemical and cellular approaches, we show how progressive truncation or phosphorylation of the GNMT N-terminus affects catalytic activity, 5mTHF-mediated inhibition, and cellular methylation potential.

## Results

### GNMT is a tissue-specific regulator of cellular SAM

While all methyltransferases utilize SAM as a methyl donor (**Figure 1A**), we asked what features may differentiate those whose activity disproportionately impacts SAM:SAH ratios. We first asked whether SAM-modulating methyltransferases possess a unique tissue distribution. We probed publicly available RNA-level datasets from the Human Protein Atlas^45^ and examined transcript abundances (measured in normalized transcript abundance in peak tissue expression) of all known human methyltransferases across tissue types. These values were plotted against a given methyltransferase’s tissue specificity score, tau^46^. This analysis revealed that SAM-modulating methyltransferases such as GNMT, PEMT, NNMT and COMT clustered with other highly tissue-specific methyltransferases (**Figure 1B**). Among these, GNMT was exclusively enriched at high levels in the liver and pancreas (**Figure 1C**).

**Figure 1.**
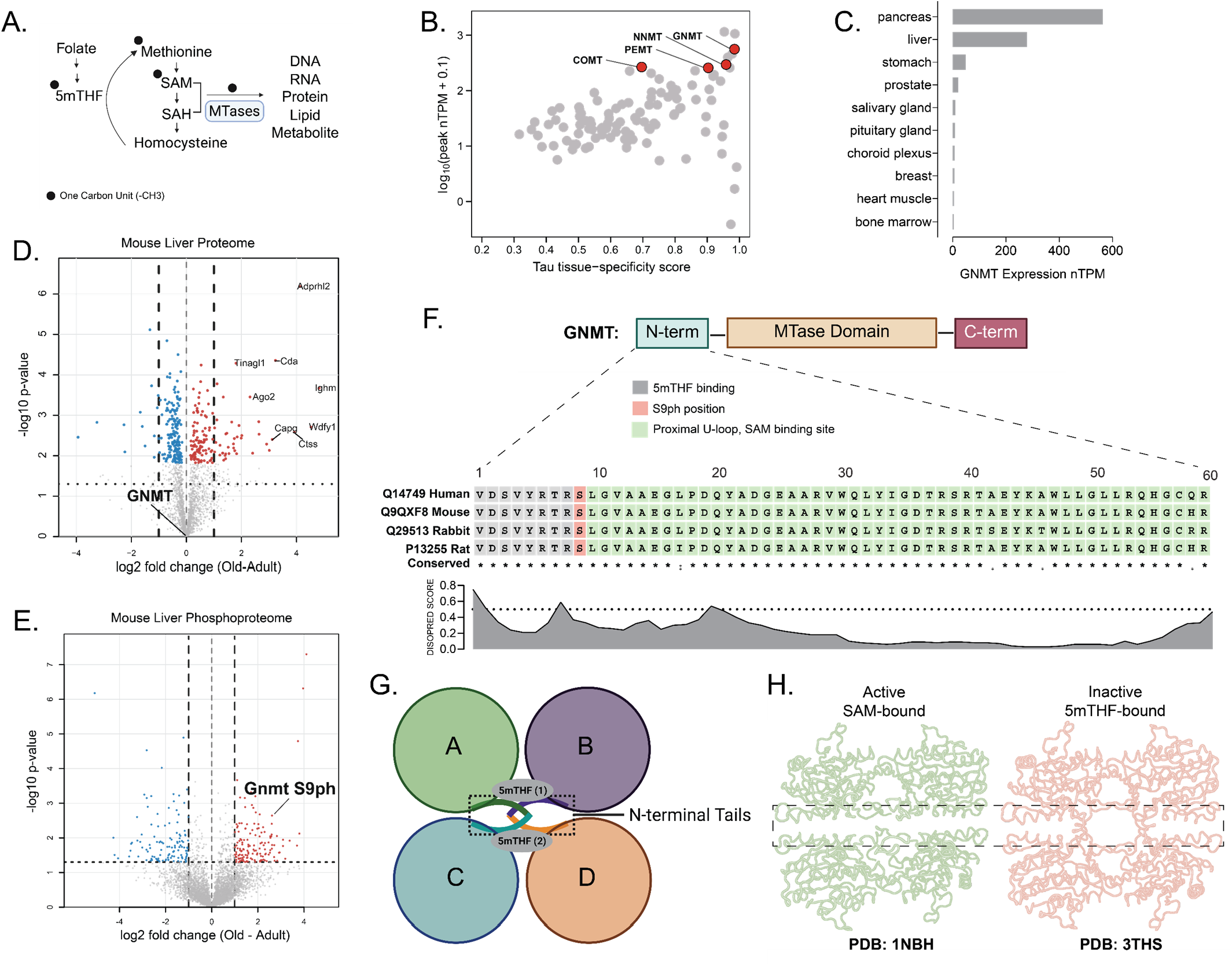
GNMT is a tissue-specific methyltransferase with age-associated N-terminal S9 phosphorylation. **(A)** Schematic of one-carbon metabolism showing folate-derived 5mTHF, methionine, SAM, SAH, homocysteine, and SAM-dependent methyltransferase reactions on DNA, RNA, protein, lipid, and metabolite substrates. **(B)** Scatter plot of human methyltransferases comparing peak tissue expression level, plotted as log10 peak nTPM + 0.1, against tissue-specificity score, tau, using Human Protein Atlas transcript abundance data. Highlighted methyltransferases include GNMT, NNMT, PEMT, and COMT. **(C)** Bar plot showing GNMT transcript abundance across human tissues. **(D)** Volcano plot of differential protein abundance in old versus adult mouse liver proteome data from Mohallem et al., with log2 fold change on the x-axis and −log10 p value on the y-axis. **(E)** Volcano plot of differential phosphorylation abundance in old versus adult mouse liver phosphoproteome data from Mohallem et al., with GNMT S9 phosphorylation labeled. **(F)** GNMT domain schematic showing the N-terminal region, methyltransferase domain, and C-terminal region, with an expanded alignment of the GNMT N-terminus across human, mouse, rabbit, and rat sequences. The alignment marks residues associated with 5mTHF binding, the S9 phosphorylation site, and the proximal U-loop/SAM-binding region; the DISOPRED conservation track is shown below. **(G)** Schematic representation of the GNMT tetramer showing chains A–D, the positioning of opposing N-terminal tails, and two bound 5mTHF molecules at the intersubunit interface. **(H)** Structural views of GNMT based on PDB: 1NBH and PDB: 3THS, with the N-terminal tail/interface region indicated by dashed boxes.

Previous work from our groups showed that GNMT expression is reduced in aged rats (19 months old) compared to young adult rats (3 months old) and that this loss of expression can be ameliorated with dietary restriction^37^. This observation suggested that GNMT expression and activity is responsive to age- and dietary restriction–dependent regulation. Since post translational modification of GNMT (i.e. phosphorylation) has previously been shown to regulate GNMT independent of protein abundance^34,47,48^, we asked whether specific phosphorylation marks may be enriched either in aged versus young animal models. We therefore examined publicly available liver phosphoproteomics datasets for age-associated phosphorylation events on GNMT. Differential proteomic and phosphoproteomic expression analysis of old versus young adult mice from Mohallem *et. al*.^49^ revealed that, while there was no difference in overall hepatic GNMT protein abundance between old and young adult mice in this study, phosphorylation on serine 9 (S9ph) was one of the most highly enriched marks in the livers of aged mice (**Figures 1D, E**). Notably, in a new phosphoproteomic study of sulfur amino acid restricted and *ad libidum* fed mice, we observed modestly lower occupancy of S9ph in the livers of sulfur amino acid restricted mice than that of *adlibidum* fed mice (**Supplementary Figure S1A, Supplementary Table S1**). S9ph also appears to be decreased in hepatitis B-related hepatocellular carcinoma, supporting a putative role in the cancer tissue context^50^ (**Supplementary Figure S1B**). S9 is positioned on the partially disordered and highly conserved N-terminal tail of GNMT and is adjacent to residues that compose the 5mTHF binding pocket (**Figure 1F**). Collectively, these results suggest that S9ph is an aging and cancer associated regulatory mark, potentially impacting 5mTHF-mediated inhibition of GNMT **(Figures 1G, H)**.

### The N-terminus supports activity and SAM stability

The N-terminus has long been recognized as a dynamic gatekeeper domain modulating entry of SAM into the active site. However, the role of post translational modifications on the N-terminus, including S9ph, is poorly understood. To better resolve N-terminal regulatory features, we performed biochemical and structural studies of GNMT. We purified, and crystallized recombinant human GNMT and solved a high-resolution structure (1.65 Å, PDB: 12YI) (**Figure 2A**). Electron density was visible starting from the first residue (N-acetylvaline) in all four monomers, indicating that much of the tail adopts a reproducible conformation despite its apparent flexibility. Residues 1-8 on the N-terminus were less ordered than the adjacent U-loop composed of residues 9-23. Thus, the structure supports a model in which the GNMT N-terminal tail is structurally defined but dynamic, positioning it to serve a regulatory role.

**Figure 2.**
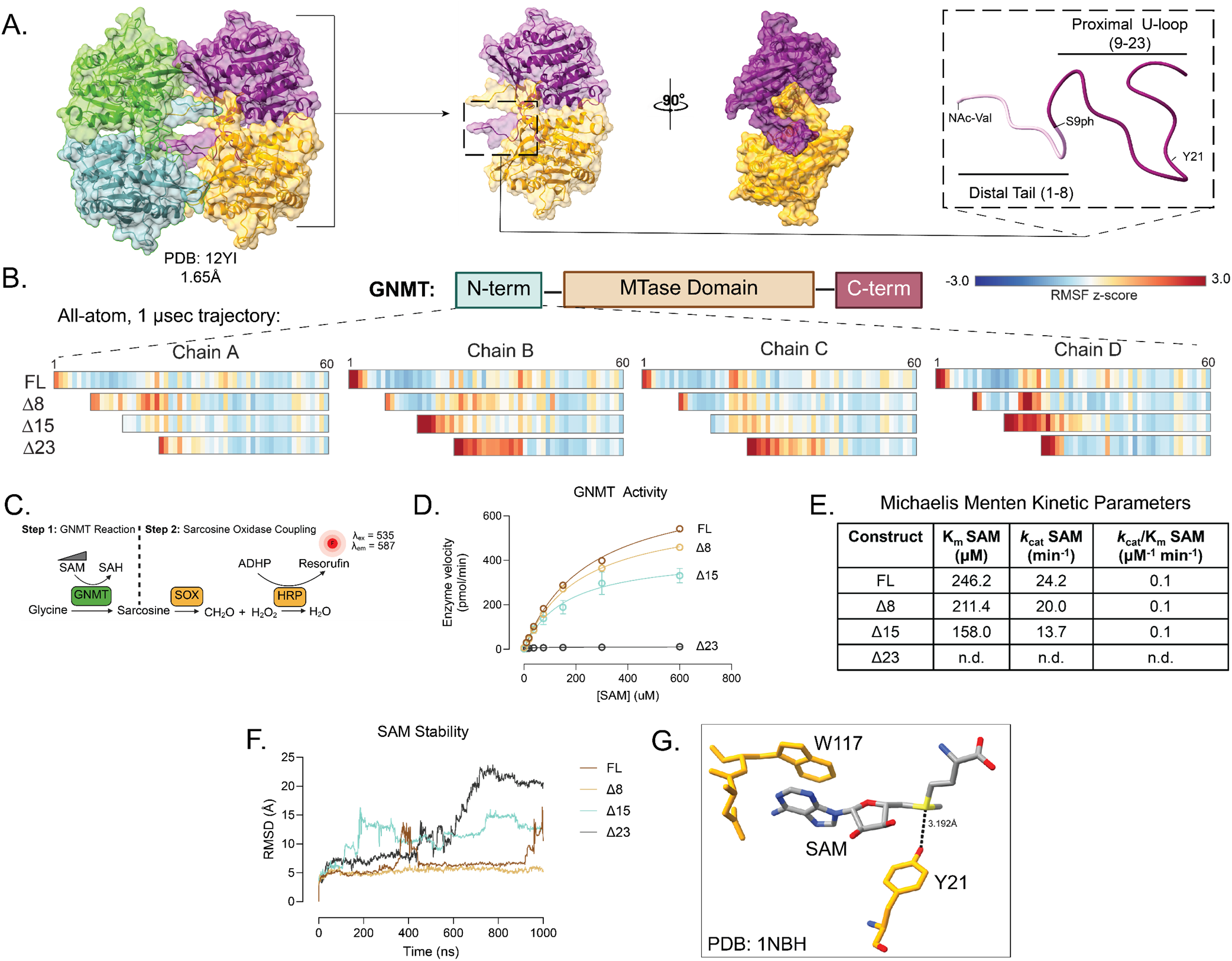
The GNMT N-terminus stabilizes the SAM-binding region and supports catalytic turnover. **(A)** Crystal structure of recombinant human GNMT solved in this study, shown as a tetramer with individual chains colored separately. The adjacent views show the GNMT dimer/interface region from two orientations, with the N-terminal region highlighted and a zoomed view of the Nterminal tail showing the distal tail residues 1–8, the U-loop residues 9–23, N-acetyl-valine, S9ph position, and Y21. **(B)** GNMT domain schematic indicating the N-terminal region, methyltransferase domain, and C-terminal region, with molecular dynamics RMSF Z-score heatmaps shown below for full-length GNMT and the Δ8, Δ15, and Δ23 N-terminal truncation variants across chains A–D. Residue positions 1–60 are shown for each chain, and the color scale indicates RMSF Z-score values. **(C)** Schematic of sarcosine oxidase–coupled enzymatic activity assay. **(D)** Sarcosine-oxidase assay used to measure GNMT activity as a function of SAM concentration for full-length GNMT, ‘Δ8, Δ15, and Δ23. **(E)** Table of kinetic parameters calculated from the activity curves in panel C, including K_m_, k_cat_, and k_cat_/k_m_ for SAM for each GNMT construct. **(F)** Molecular dynamics traces showing SAM centroid RMSD over a 1-µs simulation for full-length GNMT and the Δ8, Δ15, and Δ23 truncation variants. **(G)** Structural close-up of the GNMT SAM-binding site showing SAM positioned near W117 and Y21, with the measured distance between Y21 and SAM indicated.

We tested the dynamic features of the N-terminal tail with molecular dynamics (MD) approaches. We performed 1 microsecond all-atom MD simulations using our new high-resolution GNMT structure as the starting model. First, we investigated the impact that progressively truncated N-terminal truncation mutants might have on overall protein stability. Specific constructs used in the trajectories included full-length GNMT (FL), an 8-residue truncation mutant (Δ8), a 15-residue truncation mutant (Δ15), and a construct with complete deletion of the N-terminal tail (Δ23). For each truncation, overall tetramer stability was assessed by measuring changes in root mean square fluctuation (RMSF) across the residue index and then plotted as a heatmap of Z-score transformed data (**Figure 2B**). This analysis revealed that, with shorter N-terminal tails, overall stability of the globular domain adjacent to the tail region was compromised. Interestingly this loss of stability appeared to be chain specific, with chains B, C, and D showing the greatest degree of truncation induced instability.

We asked whether this modeled loss of globular stability might be reflected in overall enzyme activity. To test this, we generated recombinant full-length (FL), Δ8, Δ15, and Δ23 GNMT and measured enzymatic activity using a modified sarcosine oxidase–coupled assay^51,52^ **(Figure 2C)**. While V_max_ decreased with progressive truncation of the N-terminus, indicating compromised catalytic turn-over, catalytic efficiency (k_cat_/K_m_) for SAM remained comparable across FL, Δ8, and Δ15 GNMT (**Figure 2D, E**). In contrast, complete removal of the N-terminal tail (Δ23) abolished enzymatic activity. We asked whether N-terminal tail length impacts SAM stability within the active site. To address this, we performed MD simulations and measured the RMSD of the SAM centroid over a 1-microsecond trajectory. We found that SAM RMSD increased as tail length globular region of GNMT adjacent to the N-terminal tail (**Figure 2F**). Tyrosine 21 (Y21), which is also removed in the Δ23 truncation, is a critical residue for SAM binding, with its hydroxyl group forming a hydrogen bond with the sulfonium ion of SAM, thus explaining why a total loss of activity is observed with the comprehensive Δ23 truncation (**Figure 2G**)^36,53^.

As these studies showed locally induced instability adjacent to the truncated N-terminal tail, we next asked whether tail loss might more broadly disrupt GNMT tetrameric structure *in vitro*. To test this, we used analytical ultracentrifugation to measure the oligomerization states of full-length GNMT (FL) and the Δ8, Δ15, and Δ23 truncation variants across a wide concentration range, both in the decreased, consistent with reduced stability of the presence and absence of SAM (**Supplementary Figure S2**). We observed no changes in oligomerization state for any variant under the conditions tested, indicating that alterations in tail length do not detectably compromise overall tetrameric stability. Together, these data support a model in which the GNMT N-terminal tail acts as a dynamic local regulatory domain that stabilizes the SAM-binding site thereby maintaining catalytic turnover.

### The N-terminus is critical for 5mTHF feedback

GNMT is the only known human methyltransferase to be inhibited by 5-methyl-tetrahydrofolate (5mTHF), the most abundant reduced folate species in mammalian cells, which carries a one carbon unit used in re-methylation of homocysteine into methionine (**Figure 3A**). Based on prior structural models (e.g. PDB: 3THS)^54^, the distal N-terminal tail (residues 1-8) comprises a significant portion of the putative (5mTHF) binding site, thereby implicating the tail as a negative regulatory domain (**Figure 1G, H**). Closer inspection of the 5mTHF-bound structure revealed specific residues on the N-terminus, namely two Y5 side chains on antiparallel tails engaging the pterin and para-aminobenzoic (PABA) ring systems on 5mTHF (**Figure 3B**). In support of the structural observations implicating the N-terminus in 5mTHF binding, we performed MD simulations to determine how 5mTHF stability might be impacted by tail truncation. Over a 1 microsecond trajectory, RMSD at the 5mTHF centroid increased with decreasing tail length, indicating that the tail is likely an indispensable component of the 5mTHF binding pocket (**Figure 3C**). To validate these *in silico* findings, we performed isothermal calorimetry experiments to measure altered thermodynamics of 5mTHF binding with respect to tail length. These studies demonstrated that, while 5mTHF bound GNMT FL exothermically with a K_D_ of 6.25 μM (**Figure 3D, 3E**), 5mTHF failed to bind GNMT Δ8 and Δ15 (**Figure 3F, 3G**).

**Figure 3.**
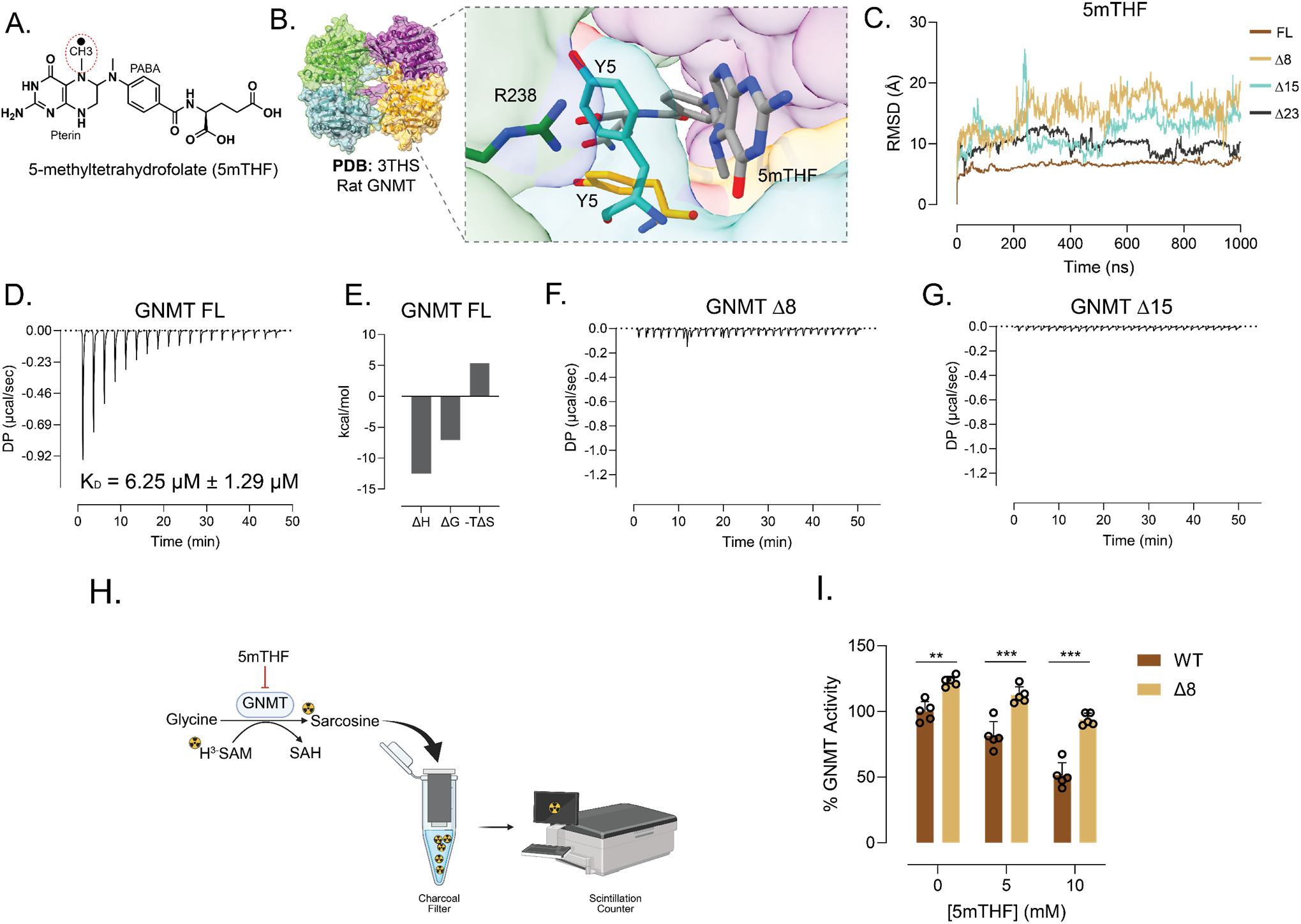
The distal GNMT N-terminal tail is required for 5mTHF binding and feedback inhibition. **(A)** Chemical structure of 5-methyltetrahydrofolate (5mTHF), with the methyl group, pterin ring, and PABA moiety labeled, corresponding to the folate metabolite. **(B)** Structure of the GNMT tetramer from PDB: 3THS with an expanded view of the 5mTHF-binding site, showing 5mTHF positioned near the N-terminal Y5 residues and the labeled R238 residue (R241 in humans). **(C)** Molecular dynamics trace showing 5mTHF RMSD over a 1-µs simulation for full-length GNMT, Δ8, Δ15, and Δ23. **(D)** Isothermal titration calorimetry thermogram for 5mTHF titration into full-length GNMT, with the fitted KD value shown. **(E)** Thermodynamic parameters from the full-length GNMT ITC experiment, including ΔH, ΔG, and −TΔS. **(F)** Isothermal titration calorimetry trace for 5mTHF titration into GNMT Δ8. **(G)** Isothermal titration calorimetry trace for 5mTHF titration into GNMT Δ15. **(H)** Schematic of the radiolabeled SAM GNMT activity assay using glycine, 3H-SAM, 5mTHF, charcoal filtration, and scintillation counting. **(I)** Quantification of GNMT activity for WT and Δ8 enzymes at the indicated 5mTHF concentrations.

To correlate the observed loss of binding to inhibitory capacity, we performed an inhibition assay using a radioactive tritiated-SAM (^3^H-SAM) substrate (**Figure 3H**). This assay revealed that, compared to GNMT FL, 5mTHF demonstrated a reduced inhibitory capacity against GNMT Δ8 (**Figure 3I**). Overall, these results indicate that the N-terminal tail is critical for maintaining negative feedback inhibition by 5mTHF.

### S9ph is poised to disrupt the 5mTHF binding pocket

As described above, the GNMT N-terminus harbors multiple post-translational modifications, including phosphorylation at serine 9 (S9ph). Structural modeling places S9 within the U-loop, adjacent to the 5mTHF-binding site (**Figure 4A**), prompting us to ask whether S9ph affects N-terminal tail dynamics and 5mTHF-mediated inhibition. To test this, we generated a GNMT S9E phosphomimetic mutant and measured 5mTHF binding thermodynamics by isothermal titration calorimetry. This assay revealed a complete loss of detectable 5mTHF binding to GNMT S9E, suggesting that S9ph may dynamically prevent 5mTHF inhibition (**Figure 4B**). Molecular dynamics simulations showed that S9ph weakens interactions between the GNMT–5mTHF complex and key residues that interact strongly in the non-phosphorylated complex (**Figure 4C**). Among the residues whose interactions were disrupted by S9ph, R241 showed one of the highest interaction frequencies with 5mTHF in the non-phosphorylated context; upon S9 phosphorylation many of these interactions were reduced across the 1-microsecond trajectory (**Figure 4D**). Consistent with our model of a disrupted 5mTHF-binding pocket upon S9 phosphorylation, distance measurements across the trajectory revealed that R241 moves away from 5mTHF in the presence of S9ph (**Figure 4E**). The same simulations indicated that R241 frequently contacts the PABA and pterin rings of 5mTHF (**Supplementary Figure S3**).

**Figure 4.**
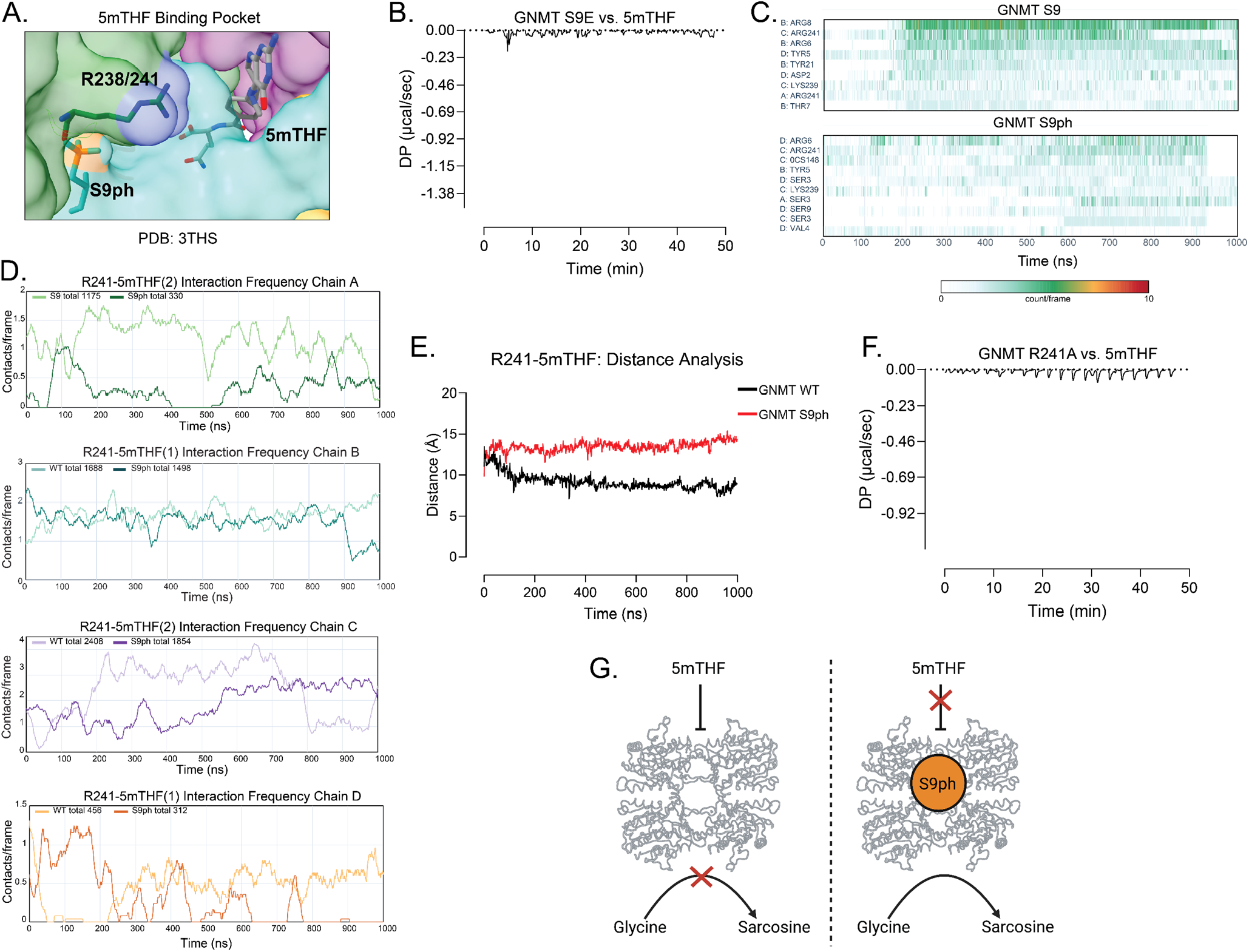
S9 phosphorylation disrupts the 5mTHF-binding pocket and relieves GNMT feedback inhibition. **(A)** Structural view of the GNMT 5mTHF-binding pocket based on PDB: 3THS, showing 5mTHF positioned near S9ph and R238 (R241 in humans). **(B)** Isothermal titration calorimetry thermogram for 5mTHF titration into GNMT S9E showing absence of isotherm. **(C)** Molecular dynamics interaction heatmaps showing residue contacts with 5mTHF over a 1-microsecond trajectory for GNMT S9 and GNMT S9ph, with interacting residues listed on the y-axis and simulation time shown on the x-axis. **(D)** Interaction-frequency traces for R241 contacts with 5mTHF across chains A–D during molecular dynamics simulations, comparing the indicated GNMT S9 and S9ph conditions. **(E)** Distance analysis measuring R241–5mTHF separation over the molecular dynamics trajectory for GNMT WT and GNMT S9ph. **(F)** Isothermal titration calorimetry thermogram for 5mTHF titration into GNMT R241A. **(G)** Model schematic summarizing 5mTHF-dependent inhibition of GNMT in the absence or presence of S9ph.

To test these computational observations, we generated a recombinant GNMT R241A mutant to disrupt the putative R241–5mTHF interaction. Binding assays showed that 5mTHF failed to bind GNMT R241A, indicating that R241 is a critical and previously unrecognized residue within the 5mTHF-binding pocket (**Figure 4F**). Taken together with the previous observation that S9ph was enriched in livers of aged mice (**Figure 1E**), these results position S9ph as a dynamic phosphoregulatory mark which modulates 5mTHF-mediated negative feedback of GNMT (**Figure 4G**).

### N-terminus maintains cell growth and SAM levels

To examine cellular consequences of GNMT activity, we overexpressed GNMT FL, Δ8, and S9E in HepG2 (hepatocellular carcinoma) and MIHA cells (immortalized non-cancerous hepatocytes), both of which lack endogenous GNMT expression (**Figure 5A)**. We first measured GNMT activity in these cells by quantifying secreted sarcosine in the cell culture media using the sarcosine oxidase assay (**Figure 5B**). Sarcosine production, non-existent in the absence of GNMT (GFP control), was significantly increased in cells expressing Δ8 relative to FL. This observation suggested that, despite having decreased velocity *in vitro*, putative loss of inhibitory capacity by 5mTHF renders a constitutively active, gain of function enzyme (**Figure 5C**). To definitively test whether this gain of function was due to loss of negative feedback by 5mTHF, we treated HepG2 cells expressing these mutants with leucovorin, a potent folate agonist, and measured sarcosine production (**Figure 5D, Supplementary Figure S4A**). We found that, while leucovorin treatment of GNMT FL-expressing cells lowered media sarcosine abundance, cells expressing GNMT Δ8 and S9E showed no response, thus demonstrating their loss of negative feedback regulation by 5mTHF.

**Figure 5.**
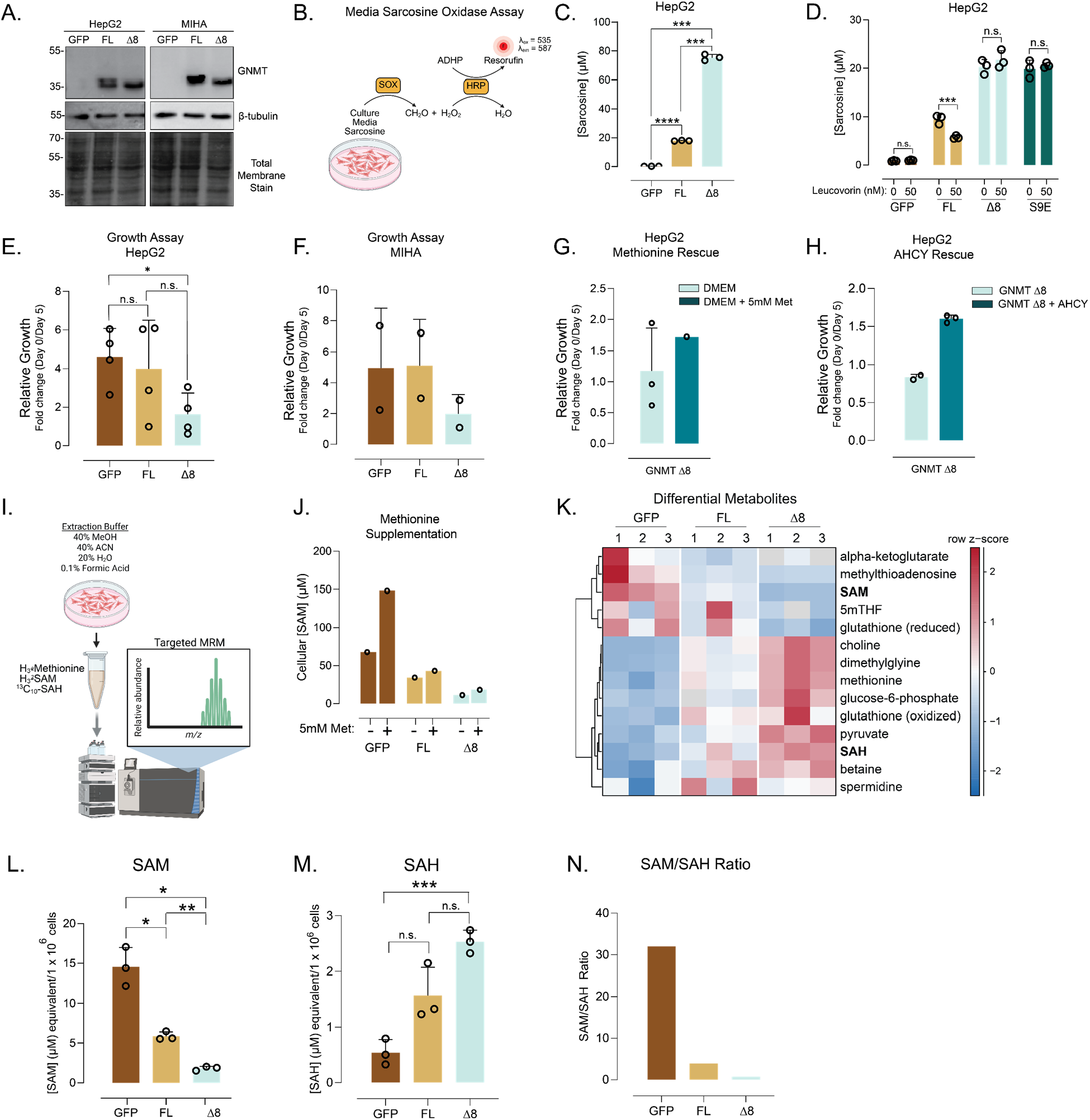
Loss of GNMT feedback control depletes SAM and impairs hepatocyte growth. **(A)** Immunoblot analysis of GNMT expression in HepG2 and MIHA cells expressing GFP control, GNMT FL, and Δ8. **(B)** Schematic of the media sarcosine oxidase assay used to quantify secreted sarcosine from cultured cells, in which sarcosine is coupled to SOX- and HRP-dependent resorufin production. **(C)** Quantification of media sarcosine concentration in HepG2 cells expressing GFP, GNMT-FL, and Δ8. **(D)** Quantification of media sarcosine concentration in HepG2 cells expressing GFP, GNMT FL, Δ8, and S9E following treatment with 0 or 50 nM leucovorin. **(E)** Relative growth of HepG2 cells expressing GFP, GNMT FL and Δ8, plotted as fold change from day 5 relative to day 0. **(F)** Relative growth of MIHA cells expressing GFP, GNMT FL, and Δ8, plotted as fold change from day 5 relative to day 0. **(G)** Relative growth of HepG2 cells expressing GNMT Δ8 cultured in DMEM or DMEM supplemented with 5 mM methionine. **(H)** Relative growth of HepG2 cells expressing GNMT Δ8 +/-AHCY overexpression (fold change = 1.5, SEM = 0.03). **(I)** Schematic of the targeted metabolomics workflow used to measure one-carbon metabolites from cultured cells by LC–MS/MS. **(J)** Cellular SAM concentration in HepG2 cells expressing GFP, GNMT FL, and Δ8 cultured with or without 5 mM methionine. **(K)** Differential metabolite abundance across GFP, GNMT FL, and Δ8 represented as a row-z-score heatmap. Absolute concentration of SAM **(L)** and SAH **(M)** across GFP, GNMT FL, and Δ8. **(N)** SAM/SAH ratios across GFP, GNMT FL, and Δ8.

Next, we asked whether a gain of function in GNMT may alter cellular phenotypes such as growth rate. Growth rate analysis revealed a proliferative defect in HepG2 and MIHA cells expressing GNMT Δ8 following 5 days in culture post transduction and antibiotic selection (**Figure 5E, F**). We hypothesized that this growth defect may result from depletion of SAM available for critical methyltransferase reactions by constitutively active GNMT Δ8. To test this, we cultured HepG2 cells expressing GNMT Δ8 with DMEM supplemented with excess methionine (5mM), as this approach has previously been shown to rescue SAM levels^14^. However, no growth rescue was observed following methionine supplementation, suggesting that the observed growth defect was independent of SAM abundance (**Figure 5G**).

Since SAH is a known product inhibitor of methyltransferases^55,56^, we asked whether SAH accumulation from constitutive GNMT activity might cause the observed growth defect. To test this, we over-expressed S-adenosylhomocysteine hydrolase (AHCY) in GNMT Δ8 expressing cells (**Supplementary Figure S4B**), which converts SAH to homocysteine, to lower SAH levels and determine whether this could rescue the defect. Notably, AHCY overexpression led to a partial rescue of the growth defect (**Figure 5H**), implicating SAH product inhibition as a key mechanism and motivating metabolomics studies to investigate dysregulation of methyl-donor homeostasis.

Targeted metabolomics (**Figure 5I**) revealed that, while, methionine supplementation substantially increased intracellular SAM concentrations in GFP-expressing control cells, this had no effect in cells expressing GNMT FL or Δ8 (**Figure 5J**). This suggests that rapid GNMT-mediated SAM turnover prevents SAM accumulation. Given that methionine is the direct metabolic precursor to SAM, we asked whether GNMT expression may increase cellular demand and uptake of methionine. To test this, we performed a cellular uptake experiment with a radioactive methionine tracer (^3^H-Met). Notably, after 1 hour of incubation with tracer, no change in radioactivity was observed in any of the conditions (**Supplementary Figure S4C, S4D**).

We measured the abundance of 14 metabolites related to the one carbon pathway in HepG2 cells, including methionine, SAM, SAH, and glutathione. (**Figure 5K**). In HepG2 cells expressing GNMT FL, SAM levels decreased by approximately 50% relative to GFP controls, whereas GNMT Δ8 expression led to an approximately 80% reduction in SAM, consistent with loss of 5mTHF-dependent feedback inhibition (**Figure 5L**). These changes were accompanied by a corresponding increase in SAH levels (**Figure 5M**). Overall, these changes contributed to a reduced SAM/SAH ratio in GNMT Δ8 (**Figure 5N**), indicating a decreased methyl donor pool and decreased methylation potential.

### GNMT N-terminus regulates methylation dynamics

To determine whether GNMT-mediated SAM depletion alters cellular protein methylation dynamics, we assessed arginine^57^ and lysine methylation^58^ (**Figure 6A**) in cells expressing GNMT FL or GNMT Δ8 by immunoblotting. GNMT Δ8 expression resulted in a marked loss of asymmetric dimethylarginine (Rme2a), consistent with reduced type I PRMT activity. We also observed a decrease in symmetric dimethylarginine (Rme2s) accompanied by an increase in monomethylarginine (Rme1) (**Figure 6B**). Lysine methylation profiles were also altered by GNMT tail truncation, with reduced trimethyl-lysine (Kme3) and increased mono-(Kme1) and dimethyl-lysine (Kme2) (**Supplementary Figure S5A**). These parallel observations were consistent with broad perturbations in methylation capacity under conditions of GNMT Δ8 mediated SAM depletion and SAH increase. Consistent with no increase in SAM levels described above, methylation levels were not rescued with methionine supplementation in cells expressing GNMT Δ8 (**Figure 6C**). To determine whether elevated SAH levels decrease arginine methylation through product inhibition of protein arginine methyltransferases (PRMTs), we overexpressed AHCY in HepG2 cells expressing GNMT Δ8 to reduce SAH levels. AHCY overexpression partially rescued the Rme2s signal, which was reduced by GNMT Δ8 expression alone, suggesting that Type II PRMTs may be sensitive to SAH-mediated product inhibition (**Supplementary Figure S5B**). Overall, these findings suggest that regulation of GNMT activity is critical for maintaining global protein methylation.

**Figure 6.**
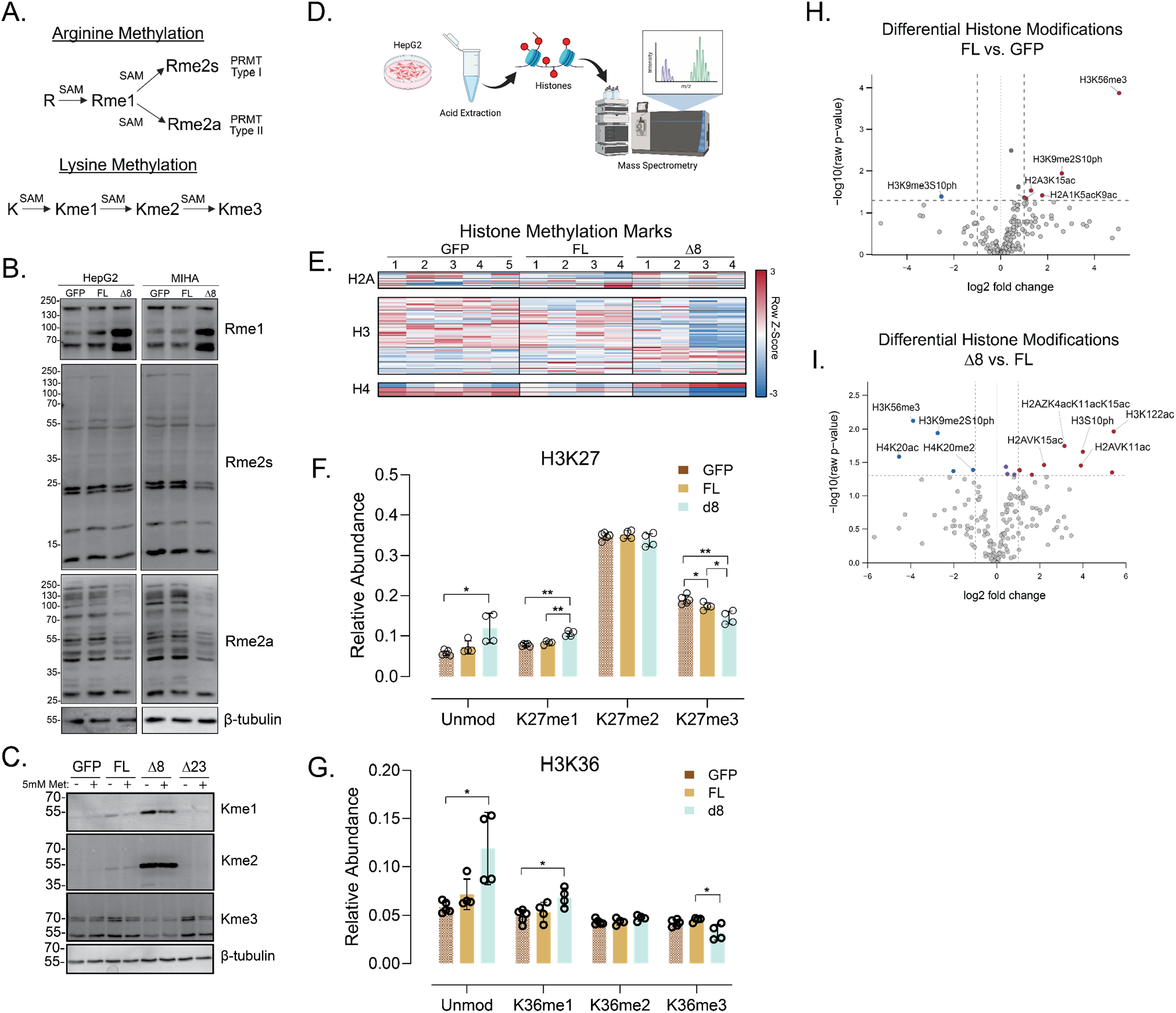
Constitutively active GNMT disrupts protein and histone methylation landscapes. **(A)** Schematic representing stepwise, SAM-dependent, enzymatic methylation of arginine and lysine. **(B)** Representative immunoblots of global monomethylarginine (Rme1), symmetric dimethylarginine (Rme2s), and asymmetric dimethylarginine (Rme2a) in cells expressing GFP control, GNMT FL, and Δ8, with loading controls. **(C)** Representative immunoblots of monomethyllysine (Kme1), dimethyllysine (Kme2), and trimethyllysine (Kme3) marks in the indicated GNMT-expressing conditions +/-supplemental (5mM) methionine, shown with total protein or loading controls. **(D)** Schematic of the histone post-translational modification workflow, showing HepG2 cell collection, acid extraction of histones, and mass spectrometry analysis. **(E)** Heatmap of relative histone methylation abundance across biological replicates from HepG2 cells expressing GFP, GNMT FL, and Δ8. **(F)** Quantification of H3K27 methylation states across GFP, GNMT FL, and Δ8 conditions. **(G)** Quantification of H3K36 methylation states across GFP, GNMT FL, and Δ8 conditions. **(H)** Volcano plot of differential histone modification abundance comparing full-length GNMT with GFP control, with selected histone marks labeled. **(I)** Volcano plot of differential histone modification abundance comparing GNMT Δ8 with GNMT FL, with selected histone marks labeled.

As histones represent a major target of methyltransferases, we asked whether histone methylation marks might be altered with GNMT-mediated SAM depletion. To test this, we extracted histones from HepG2 cells expressing GFP, GNMT FL, and GNMT Δ8, digested with trypsin, and performed mass spectrometry to quantify modifications (**Figure 6D**). This analysis revealed a complex pattern of methyl-specific marks changing bi-directionally on histones H2A, H3, and H4 across conditions (**Figure 6E**). Relative abundance of H3K27me3, a repressive heterochromatic mark associated with gene silencing, was notably decreased in GNMT Δ8, while H3K24me1 was increased consistent with our previously mentioned global arginine and lysine methylation findings (**Figure 6F**). Similarly, relative abundance of H3K36me3, a mark associated with transcriptional activation, was also decreased in GNMT Δ8, while H3K36me1 was increased (**Figure 6G**). When comparing all differentially abundant marks (not exclusively methyl-marks) between GNMT FL and GFP, relatively few marks were enriched in either condition (**Figure 6H**). Among these, H3K56me3, a heterochromatic mark often co-occurring in both regulation and function with H3K9me3 was highly enriched in GNMT FL expressing cells^59^. In contrast, comparison of differentially abundant marks between cells expressing either GNMT Δ8 or GNMT FL revealed many more changes to histone modifications (**Figure 6I**). Compared with GNMT FL, GNMT Δ8 markedly reduced H3K56me3 abundance, despite H3K56me3 being strongly enriched in GNMT FL relative to GFP. Several lysine-acetyl marks were also enriched in the GNMT Δ8 condition. Overall, these results indicate that 5mTHF-mediated negative feedback through the GNMT N-terminal tail is required to maintain normal histone post-translational modification patterns. Loss of this inhibitory regulation, as modeled by GNMT Δ8, broadly remodels the histone modification landscape, including both methylation and acetylation marks.

### GNMT dysregulation activates cell stress genes

Given that loss of 5mTHF-mediated negative feedback of GNMT impaired growth, disrupted SAM homeostasis, and altered methylation state, we next asked whether these phenotypes were associated with underlying transcriptional changes. To test this, we performed 3′-end bulk RNA-seq on MIHA and HepG2 cells expressing GFP, GNMT FL, or GNMT Δ8 after 5 days of growth following puromycin selection. Principal component analysis of variance-stabilized counts showed clear separation of the three expression states in both MIHA and HepG2 cells, indicating that GNMT Δ8 induces a transcriptional program distinct from both GFP and GNMT FL, whereas GNMT FL remains closer to the control condition (**Figure 7A**). In HepG2 cells PC1 and PC2 explained 73% and 14%, while in MIHA cells, PC1 and PC2 explained 74% and 17% of the variance, respectively (**Supplementary Figure S6A**), supporting a strong and reproducible Δ8-specific effect in both models. Consistent with the relatively limited phenotypic effect of full-length GNMT, the GNMT FL versus GFP comparison produced only modest transcriptional changes in both cell lines.

**Figure 7.**
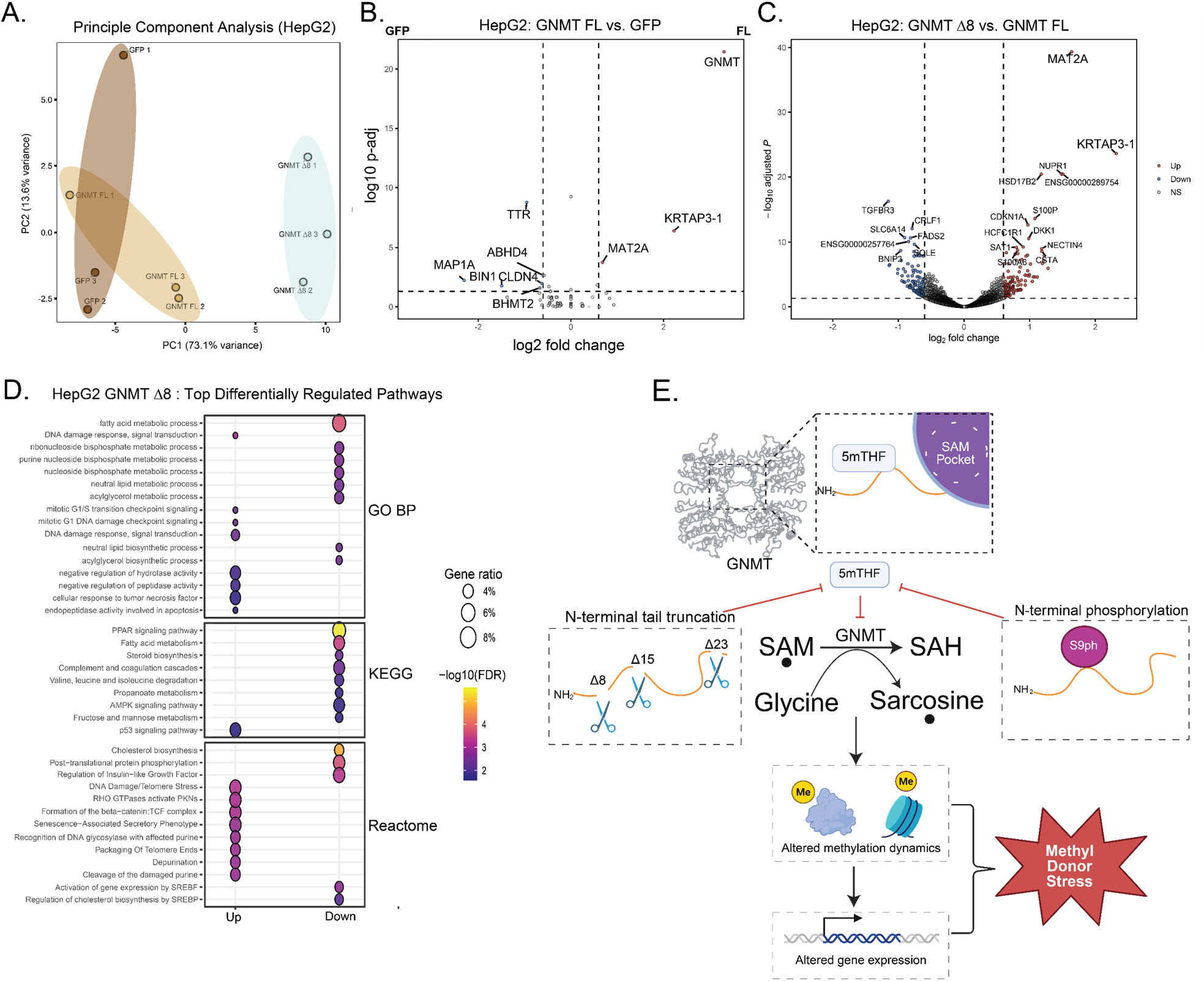
Folate-uncoupled GNMT activity induces methyl-donor stress and transcriptional remodeling. **(A)** Principal component analysis of variance-stabilized RNA-seq counts from HepG2 cells expressing GFP control, full-length GNMT, or GNMT Δ8. **(B)** Volcano plot of differential gene expression in HepG2 cells comparing full-length GNMT with GFP control, with log2 fold change on the x-axis and −log10 adjusted *P* value on the y-axis. **(C)** Volcano plot of differential gene expression in HepG2 cells comparing GNMT Δ8 with full-length GNMT, with upregulated, downregulated, and non-significant genes indicated and selected genes labeled. **(D)** Dot plot of pathway enrichment analysis from the HepG2 GNMT Δ8 versus full-length GNMT comparison, grouped by GO Biological Process, KEGG, and Reactome categories; dot size indicates gene ratio and color indicates −log10 FDR. **(E)** Summary schematic of the proposed GNMT regulatory model, showing the GNMT tetramer, the N-terminal/5mTHF/SAM-pocket region, N-terminal truncation and S9 phosphorylation, GNMT-mediated conversion of glycine to sarcosine, and associated changes in methylation dynamics, gene expression, and methyl-donor stress.

In HepG2 cells, GNMT was robustly induced, confirming successful expression, but only a small set of additional genes were upregulated including MAT2A, the enzyme that catalyzes the ATP-dependent synthesis of SAM from methionine (**Figure 7B**). MIHA cells showed a similar pattern, with strong GNMT induction and comparatively modest accompanying changes, which again included MAT2A (**Supplementary Figure S6B**).

In contrast, GNMT Δ8 expression in HepG2 cells triggered a broad transcriptional response in genes linked to one-carbon metabolism (464 up-regulated genes and 555 downregulated genes, p-adj < 0.05). GNMT Δ8 increased expression of one-carbon metabolism genes such as MAT2A, TYMS, and AHCY, while decreasing expression of BHMT2, MTR, CHDH, and MTAP, indicating that constitutively active GNMT reprograms one-carbon metabolism in association with methyl-donor stress. Genes such as SAT1 and AMD1, which are involved in polyamine synthesis and represent an alternative route of SAM utilization that by-passes SAH production, were also upregulated. Beyond one carbon metabolism-related genes, GNMT Δ8 also induced stress-response genes such as NUPR1, CDKN1A, SESN2, and EGR1.

Pathway enrichment analysis in HepG2 cells revealed distinct biological programs associated with genes up-and downregulated in GNMT Δ8 cells (**Figure 7D**). Upregulated pathways related to DNA damage response, cell-cycle checkpoint signaling, mitotic G1/S transition, p53 signaling, TNF response, and apoptotic regulation, consistent with activation of stress-response and growth regulation. Upregulated Reactome terms included DNA damage/telomere stress, RHO GTPase signaling, senescence-associated secretory phenotype, and depurination/repair-related processes. In contrast, downregulated terms in GNMT Δ8 were strongly associated with metabolic and biosynthetic pathways, including fatty acid metabolism, ribonucleotide and purine metabolism, phospholipid and acylglycerol metabolism, neutral lipid biosynthesis, and PPAR signaling. Overall, this analysis indicates that GNMT Δ8 expression is associated with a shift away from biosynthetic and metabolic processes and towards stress-response pathways leading to senescence.

Compared with the HepG2 dataset, the MIHA pathway profile revealed a distinct transcriptional response to GNMT Δ8 (**Supplementary Figure S6C**). MIHA upregulated pathways were less centered on canonical DNA damage and checkpoint pathways and instead showed enrichment for hypoxia and oxygen-response programs, reactive oxygen species metabolism, and mitochondrial membrane permeability. Both cell types showed suppression of metabolic pathways among genes down-regulated in GNMT Δ8, particularly lipid and sterol-related processes. However, the MIHA response appeared more broadly metabolic, with prominent down-regulation of fatty acid degradation, fatty acid metabolism, PPAR signaling, peroxisome, carbohydrate metabolism, nucleotide sugar biosynthesis, and cholesterol biosynthesis, whereas the HepG2 dataset more strongly emphasized lipid, nucleotide, phospholipid, and SREBP/cholesterol regulatory pathways. Reactome analysis showed enrichment of senescence-associated and IGF-related pathways across both cell lines, but these stress-associated pathways were more pronounced in the HepG2 cells than in the MIHA profile. Together, these results indicate that GNMT Δ8 induces a cell stress response in both HepG2 and MIHA cells, however the origins of that stress differ. Constitutively active GNMT expression upregulated DNA damage, check-point, and senescence programs in HepG2 cells, whereas in MIHA cells it upregulated a mitochondrial stress response.

Overall, our studies demonstrated that the GNMT N-terminus is a folate-sensing regulatory domain that couples 5mTHF-dependent feedback inhibition to control of methyl donor homeostasis (**Figure 7E**). The distal N-terminal tail supports 5mTHF binding and restrains GNMT activity, while S9ph destabilizes the inhibitory pocket and shifts GNMT toward a feedback-resistant, gain of function state. Loss of this regulation depletes SAM, elevates SAH, and compromises cellular methylation potential. These metabolic changes disrupt protein and histone methylation dynamics, impair cell growth, and activate transcriptional programs consistent with methyl-donor stress.

## Discussion

In this work, we establish the N-terminal tail of GNMT as a key regulatory domain that maintains SAM-binding site stability and coordinates 5mTHF-mediated negative feedback, thereby preserving methyl-donor homeostasis and cellular methylation potential. This mechanism is especially important given our observation that, unlike most broadly expressed methyltransferases, GNMT belongs to a subset of small-molecule methyltransferases with highly specific tissue expression and context-dependent regulation across cancer^23,60^ and aging^37,49^. This specific and context-dependent expression pattern suggests that GNMT is not merely a SAM-consuming enzyme, but a highly controlled regulator of cellular methylation potential. We showed that the distal N terminus (residues 1-8) is essential for 5mTHF-dependent inhibition and that disruption of this region uncouples catalysis from feedback control. The result is a constitutively active enzyme state that depletes SAM, elevates SAH, disrupts methylation, impairs proliferation, and induces broad stress-responsive transcriptional programs. Together, these findings provide a mechanistic explanation for how GNMT regulates cellular methylation potential and why loss of its regulation may have severe consequences for tissue homeostasis.

Our findings expand prior models of the GNMT N-terminal tail as a regulatory domain. Molecular dynamics simulations and biochemical assays showed that the N-terminal region not only acts as a gatekeeper for SAM access but also stabilizes the adjacent globular domain containing the SAM-binding site. Our results suggest that the N-terminal tail, including more distal residues that do not appear to contact SAM directly, contributes to maintaining the structural integrity and catalytic competence of the active site.

We also reinforce the previous observation^34,54^ that GNMT is negatively regulated by 5mTHF through its N-terminal tail. We identified R241 and S9 as critical residues required to maintain this inhibitory mechanism, the latter of which is preferentially phosphorylated in livers of old mice^49^, thus implicating it as a phosphoregulatory switch. We show that S9ph is modeled to disrupt residue interactions with 5mTHF, thereby destabilizing the 5mTHF-binding pocket, a model supported experimentally by the loss of detectable 5mTHF binding to GNMT S9E. Prior work has implicated several putative kinases, including PKA, in GNMT S9 phosphorylation^48,61^; however, additional studies are needed to define the signaling mechanisms that regulate this modification.

We also demonstrate the importance of 5mTHF-mediated regulation of GNMT in the cellular context, where expression of constitutively active GNMT leads to a growth defect and large perturbations to SAM:SAH balance. We demonstrate that this metabolic dysregulation is associated with loss of arginine and lysine methylation on proteins and disruption to post translational modifications on histones and transcriptional changes. These transcriptional changes suggest that constitutively active GNMT drives a reprogramming of one-carbon metabolism in response to methyl-donor stress. In both HepG2 and MIHA cells, GNMT Δ8 increased expression of genes involved in SAM synthesis, nucleotide metabolism, and SAH turnover, including MAT2A, TYMS, and AHCY, consistent with a compensatory attempt to restore methyl-donor balance. At the same time, GNMT Δ8 reduced expression of BHMT2, MTR, CHDH, and MTAP, indicating disruption of pathways that normally support methionine regeneration, choline-dependent methyl donation, and methionine salvage. This transcriptional response suggests that cells sense GNMT-driven SAM depletion but are unable to fully restore one-carbon homeostasis when GNMT activity is uncoupled from 5mTHF-dependent feedback inhibition. Thus, loss of N-terminal GNMT regulation creates a persistent methyl-donor stress state that reshapes metabolic gene expression and likely contributes to the growth defects and broader stress-response programs observed in GNMT Δ8-expressing cells.

Perturbations to SAM:SAH ratio is also connected to growth signaling through SAMTOR-dependent regulation of mTORC1^21^. SAMTOR directly senses SAM, and SAM depletion inhibits mTORC1 signaling, providing a parallel mechanism by which altered methyl-donor metabolism could limit growth.

A diverse intracellular pool of folate polyglutamates has been reported across mammalian cell types, including species that vary by one-carbon state and glutamate chain length^62,63^. The discrepancy between 5mTHF binding affinity and inhibitory capacity (see Figure 2 above) can be explained by the requirement that 5mTHF be polyglutamylated and that the N-terminal valine be acetylated to enable physiologically meaningful inhibition^34,64^. Although GNMT inhibition is sensitive to folate polyglutamylation, how the naturally mixed intracellular folate pool shapes GNMT inhibition in cells remains unresolved. Furthermore, the precise structural contribution of the polyglutamate tail to the inhibitory capacity of 5mTHF is still unclear, since density around the polyglutamate tail was absent from prior structural models (i.e. PDB: 3THS). Due to limited commercial availability and technically challenging synthesis, our *in vitro* 5mTHF studies were restricted to monoglutamylated 5mTHF, which has substantially reduced inhibitory potency against GNMT. To address this limitation and better approximate physiological interactions, we used pentaglutamylated 5mTHF in all molecular dynamics simulations. In support of a role for polyglutamylation state in GNMT regulation, cells expressing GNMT Δ8 showed altered expression of FPGS, which adds γ-linked glutamates to THF derivatives, and GGH, which hydrolyzes folate polyglutamate tails. These changes suggest that 5mTHF-mediated negative feedback may be regulated dynamically through remodeling of intracellular folate polyglutamylation states.

Prior studies have implicated S9ph as critical for the nuclear translocation of GNMT under conditions of cellular stress^48,65^, and other studies have shown that GNMT may play a non-enzymatic role in the nucleus, potentially acting as a tumor suppressor in hepatocellular carcinoma^60^. Additional studies in murine hepatocytes have demonstrated that, following chemically induced stress, nuclear translocation of GNMT is associated with engagement of the Nrf2 promoter, implicating GNMT in the regulation of this master transcription factor of redox homeostasis^66^. The mechanisms underlying this potential non-catalytic function remain poorly understood. Future work will address these open questions by combining catalytically inactive GNMT mutants (e.g. N140S) with proximity labeling and immunofluorescence approaches to define how GNMT localization and compartment-specific behavior changes in response to stimuli such as oxidative stress.

In our metabolomics studies, we observed a shift in glutathione redox balance toward oxidized glutathione (GSSG) and away from reduced glutathione (GSH), consistent with an altered cellular redox state. The interconversion between GSH and GSSG is coupled to NADPH availability, as NADPH provides the reducing equivalents to regenerate GSH from GSSG. This may be relevant to GNMT regulation because 5mTHF production requires NADPH-dependent MTHFR activity, suggesting that changes in cellular redox metabolism could influence GNMT through altered 5mTHF availability. GNMT phosphorylation may therefore provide an additional regulatory mechanism, coupling redox-dependent folate metabolism to methyl-donor consumption.

Overall, our findings position GNMT as a dynamically regulated and folate-responsive rheostat whose activity must be precisely tuned to preserve methyl donor homeostasis.

## Materials and Methods

### Expression and Purification of hsGNMT

*hs*GNMT and associated mutants were cloned into a pET29a expression vector containing a C-terminal 6x-His-tag in frame. Expression vectors were transformed into Rosetta 2(DE3) competent cells, which were expanded into a starter culture overnight at 37°C. Sterilized Luria (Miller) broth supplemented with 50 μ/mL kanamycin and 25 μ/mL chloramphenicol was inoculated with starter culture and incubated with rotation at 37°C for several hours. Once an optical density (O.D.) of 0.6 was reached, cultures were immediately induced with 1mM IPTG and incubated with rotation overnight (12-16 hours) at 18°C. The bacteria were pelleted by centrifugation, resuspended in ice cold lysis buffer (100mM Tris-HCl pH 8.0, 300 mM NaCl, 14.7 mM β-ME), and lysed by Emulsiflex (15,000-18,000 psi). *hs*GNMT was initially purified on an Ni-NTA resin (Thermo). Briefly, pre-equilibrated Ni-NTA resin was incubated with lysate at 4°C with gentle rotation for 1 hour. Beads were washed with 5 column volumes (CV) of lysis buffer supplemented with 30 mM imidazole followed by 5 CV of lysis buffer supplemented with 50 mM imidazole. Protein was eluted twice with 1.5 CV of lysis buffer supplemented with 250 mM imidazole. Purified *hs*GNMT was buffer exchanged into low salt buffer (50 mM Tris-HCl pH 8.0, 50 mM NaCl, 14.5 mM β-ME) using a BioGel P6 resin. Protein was further purified by anion exchange chromatography using a HiRes CaptoQ column (Cytiva) with a 40 CV gradient up to 1M NaCl. Pure *hs*GNMT was stored at -20°C in storage buffer (50mM Tris-HCl pH 8.0, 150-200 mM NaCl, 14.7 mM β-ME, and 50% glycerol).

### Crystallization of Apo GNMT

Midwest Center for Structural Genomics (MCSG) crystallization suites 1 and 2 (Anatrace) were used to screen for conditions for the crystallization of apo GNMT. Crystallization was performed using the sitting drop vapor diffusion method at 18 °C. 96-well INTELLI plates (Art Robbins) and the Gryphon crystallization robot (Art Robbins) were used to setup the crystallization drops, which contained 0.5 μL of [6 mg/mL] GNMT (in 50 mM Tris-HCl (pH 8), 50 mM NaCl, 2.5% glycerol, and 1 mM DTT) and 0.5 μL well solution. The volume of the well solution was 70 μL. Many conditions yielded crystals within 3 hours; however, they were too fragile to pick, so the crystals were incubated at 18 °C and grown for an additional 21 days. The well solution that yielded the best diffracting crystal was comprised of 0.2 M Ammonium formate and 20% (w/v) PEG 3350. X-ray diffraction data were collected at beamline LRL-CAT (Argonne National Laboratory, Argonne, IL) using 0.9793 Å wavelength. Diffraction data was processed using iMOSFLM^67^ and scaled with AIMLESS from the CCP4 suite^68^. SFCHECK2^68^ and XTRIAGE3^69^ were used to analyze the data quality. Matthews coefficients (Vm) were used to estimate the number of monomer molecules in the unit cell. GNMT crystal structures were determined by molecular replacement using PHASER^70^. GNMT (Protein Data Bank (PDB) entry 1R74) chain A was used as the initial phasing template. COOT^71^ was used to manually adjust and complete the model obtained from PHASER. Refinements were performed with REFMAC5, using standard protocols for NCS refinement^72^. All structural measurements and molecular graphics were generated using the UCSF Chimera package from the Computer Graphics Laboratory, University of California, San Francisco (supported by National Institutes of Health Grant P41 RR-01081)^73^.

### Sarcosine assay to measure GNMT activity

We used a sarcosine oxidase (SOX)-coupled enzyme assay^51^ to measure enzyme activity of *hs*GNMT and associated mutants. SOX from *bacillus* sp. (Uniprot accession: P40859) cloned into a pET28a(+) vector and expressed in Rosetta2 (DE3) E. coli. SOX was purified by Ni-NTA affinity chromatography as described previously^74^. SAM was titrated in the presence of 312 nM *hs*GNMT and 20mM glycine (100 μL total reaction volume in 50 mM Tris pH 8.0 buffer). The reaction mixture was incubated at 30 °C for up to 10 minutes and quenched via the addition of 30 μL of 1 M acetic acid. The pH of the reaction mixture was adjusted with 30 μL of 1 M Tris base prior to proceeding to the SOX reaction. 50 μL of the reaction mixture (and a serial dilution of sarcosine as a standard curve) was combined with 50 μL of a mastermix containing 4 mU/mL of SOX, 10 mU/mL of horse-radish peroxidase (HRP), and 1 mM of 10-Acetyl-3,7-dihydroxyphenoxazine (ADHP/Amplex Red) and incubated at 37°C in the dark for 30 minutes. Absorbance at 570 nm (or fluorometry at λ_em_ = 535 nm and λ_ex_ = 587 nm) was measured using a SpectraMax-iD5 plate reader.

### Isothermal Calorimetry (ITC)

ITC experiments were performed on the Malvern MicroCal PEAQ. For all ITC experiments, *hs*GNMT was buffer exchanged into ITC buffer (50 mM HEPES pH 8.0, 150 mM NaCl, 1 mM TCEP) and concentrated to 30-60 μM. Ligand (i.e. 5mTHF, SAM) solutions were prepared in ITC buffer at concentrations ranging from 1-3 mM. Each experiment included 19-25 ligand injections ranging from 1-3 μL each. Data was processed and analyzed using Malvern MicroCal PEAQ software. Isotherm curves were assigned based on the best fit.

### Analytical Ultracentrifugation

Analytical ultracentrifugation (AUC) studies were conducted of wild type or truncated GNMT (9.3 μM, monomer) in 150 mM NaCl, 50 mM HEPES, pH 8.0, and 1mM TCEP in the absence or presence of 0.5 or 1.0 mM SAM. The experiments were conducted in the AN-60 Ti rotor of a Beckman XL-I analytical ultracentrifuge using cells assemble with double sector charcoal EPON centerpiece and sapphire glass windows. Centrifuge runs were conducted at 48,000 rpm with sedimentation monitored at a detection wavelength of 290nm at 4 °C. The samples were equilibrated in the centrifuge for 3 hr. Sedimentation scans were collected the sample had completely sedimented to the base of the cell. A c(s) component analysis was conducted using the program Sedfit version 17.0 to deconvolute the species present in a solution^75^. The c(s) species distributions were exported to the graphing program Prism to generate the distributions shown in the text and the supporting information.

### Protein/Ligand Preparation and Molecular Docking

The apo human GNMT crystal structure (resolution: 1.65 Å) was processed for its topology refinement, energy minimization, and side-chain fixation using the Protein Preparation panel of Maestro (Schrödinger 2023-2). S-adenosyl-L-methionine (SAM) and 5-methyltetrahydrofolate pentaglutamate (5mTHF-Glu_5_) were prepared using LigPrep (Schrödinger 2023-2). Their protonation, stereochemical and tautomeric states were assigned using Epik at pH 7.0 ± 2.0, and structures were energy-minimized with the OPLS3 force field prior to docking. Four SAM structures and two 5THF-Glu_5_ structures were docked into the GNMT structure via induced fit docking (Schrödinger 2023-2), incorporating receptor flexibility through softened van der Waals radii, Prime refinement and Glide redocking. The docked poses of SAM and 5mTHF-Glu_5_ were selected based on their respective binding poses in the RCSB PDB structures (PDB: 1NBH and 3THS) and IFD scores.

### Molecular dynamics simulations

The low-energy 3D structures of apo GNMT, GNMT-SAM-5mTHF-Glu_5_ complex, GNMT Δ8-SAM-5mTHF-glu_5_ complex, GNMT Δ15-SAM-5mTHF-glu_5_ complex, GNMT Δ23-SAM-5mTHF-glu_5_ complex, GNMT S9ph-SAM-5mTHF-glu_5_ complex, GNMT acV1-SAM-5mTHF-glu_5_ complex, and GNMT R241A-SAM-5mTHF-glu_5_ complex were submitted to Desmond System Builder (Schrödinger 2023-2), where a predefined TIP3P solvent model was applied within a volume-minimized orthorhombic box. The system was neutralized with Na^+^ ions, and 0.15 M NaCl was added under the OPLS4 force field. As a result, the apo GNMT model system contained 118189 atoms; GNMT-SAM-5mTHF-glu_5_ contained 118408 atoms; GNMT Δ8-SAM-5mTHF-glu_5_ model system contained 118175 atoms; GNMT Δ15-SAM-5mTHF-glu_5_ model system contained 118041 atoms; GNMT Δ23-SAM-5mTHF-glu_5_ model system contained 118214 atoms; GNMT S9ph-SAM-5mTHF-glu_5_ model system contained 117331 atoms; GNMT acV1-SAM-5mTHF-glu_5_ model system contained 118407 atoms; while GNMT R241A-SAM-5mTHF-glu_5_ model system contained 118251 atoms. The prepared Desmond model systems were then loaded onto Molecular Dynamics (MD) panel (Schrödinger 2023-2) to run a 1000-ns all-atom MD simulation with 1.0-ps recording interval. Each system was first relaxed before the production run, which was performed under isothermal-isobaric (NPT) ensemble conditions. The Nose– Hoover Chain thermostat (1.0 ps of relaxation time) and Martyna–Tobias–Klein barostat (2.0 ps of relaxation time) were used to maintain the temperature and pressure, respectively. MD simulations were performed 3 times for each model system using different random seed numbers to ensure statistical reliability. RMSD and RMSF were computed for the protein backbone and ligands (SAM and 5mTHF-glu_5_) over the trajectory phase of the MD simulations on Maestro, in which the structures were aligned to their respective initial frame. Distances were measured between the Cα atoms of the indicated residues across all trajectory frames on Maestro.

### Cell culture

HEK293T (ATCC: CRL-3216), HepG2 (ATCC: HB-8065), and MIHA (Montefiore-Einstein Liver Center) cells were cultured in DMEM with 10% FBS and 1% penicillin/streptomycin. Transient overexpression of *hs*GNMT and associated mutants was achieved with calcium phosphate transfection of the corresponding pcDNA3.3 mammalian overexpression plasmid, which was modified to include an internal ribosome entry site (IRES) downstream of the open reading frame to allow for bicistronic expression of GFP as a fluorescent marker of overexpression. Media was changed 8 hours after transfection and degree of overexpression was assessed the following day by fluorescent microscopy and immunoblotting. For stable overexpression of GNMT and associated mutants, lentivirus was generated using a calcium phosphate precipitation method. Briefly, proteins of interest were cloned into the mammalian overexpression vector pLEX307 using a Gateway cloning strategy. The target pLEX307 plasmid was mixed with lentiviral packaging plasmids at a 1:1:1 molar ratio in HEPES-buffered saline and calcium chloride. Cells were transduced with lentivirus by centrifugation (1000 × g, 37 °C, 1.5 h) in the presence of 10 µg/mL polybrene. The following day, media was replaced with DMEM containing 5 µg/mL puromycin for selection. After 2 days of selection, cells were replated for downstream experiments. Successful selection was confirmed by assessing the viability of non-transduced control cells. For cell culture experiments involving leucovorin, DMEM deficient in folates and other relevant metabolites was purchased from the Memorial Sloan Kettering Cancer Center Media Preparation Core.

### Media sarcosine quantification

Sarcosine in the media was quantified using the SOX-coupled assay described above.

### Methionine uptake assay

Five days after puromycin selection, cellular methionine uptake was measured by incubating cells in DMEM containing 5 million CPM of tritiated methionine (^3^H-met) for 1 hour at 37 °C. Cells were then washed twice with PBS and serially scraped three times with 0.5 mL PBS. The collected suspension was added directly to 10 mL of scintillation fluid and radioactivity was measured using a scintillation counter (Perkin Elmer).

### Targeted Metabolomics

Metabolites from cultured cells were extracted using an ice-cold extraction buffer composed of 40% acetonitrile, 40% methanol, 20% water, and 0.1% formic acid, supplemented with stable isotope-labeled SAM, SAH, glycine, and sarcosine as internal standards for spike-in normalization. The cell–extraction buffer mixture was scraped into microcentrifuge tubes, and insoluble debris was pelleted by centrifugation (18,000 × g, 4 °C, 10 min). The supernatant was transferred to fresh tubes, flash-frozen in liquid nitrogen, and dried overnight in a SpeedVac. Dried metabolites were resuspended in 0.1% formic acid, clarified by centrifugation, and 8 microliters were injected into an LC–MS Agilent 6490 triple quadrupole (QQQ) system equipped with a Phenomenex Luna 100 Å 75 × 2.0 mm C18 column. A multi-reaction monitoring (MRM) approach was used to quantify dozens of metabolites (see **Supplementary Table S2** for MRM parameters). The mobile-phase gradient was delivered at a flow rate of 0.4 mL/min over a 13.01-minute run. The LC method began at 100% solvent A (0.1% formic acid) and 0% solvent B (95% acetonitrile, which was maintained until 2.00 min. Solvent B was then increased to 100% by 7.00 min and held at 100% B until 9.50 min. The gradient was returned to initial conditions of 100% A and 0% B at 9.51 min and maintained through 13.00 min for column re-equilibration. At 13.01 min, the method switched to 0% A and 100% B.

### Immunoblotting

Cells were washed with ice-cold PBS and lysed in RIPA buffer supplemented with 1X HALT protease inhibitor (Thermo). Lysates were incubated on ice for 30 min and clarified by centrifugation at 10,000 × g for 10 min at 4°C. Protein concentrations were measured using a BCA protein assay. Equal amounts of protein were denatured in Laemmli sample buffer with reducing agent at 95°C for 5 min, separated by SDS– PAGE, and transferred to PVDF membranes. Membranes were blocked for 1 h at room temperature in 1% ECL Advance Blocking Reagent (Cytiva) in PBS with 0.01% Tween20 and then incubated overnight at 4°C with primary antibodies diluted in PBS supplemented with 5% BSA, 0.05% sodium azide, and 0.01% Tween20. The following primary antibody dilutions were used: GNMT 1:1000 (ProteinTech 18790-1-AP), β-tubulin 1:50,000 (DSHB E7), GFP 1:5000 (ProteinTech50430-2-AP), Rme1 1:2000 (Cell Signaling Technologies 13522S), Rme2s 1:10,000 (provided by Mark Bedford, MD Anderson), Rme2a 1:10,000 (Cell Signaling Technologies 8015S), Kme1 1:10,000 (CST 14680), Kme2 1:10,000 (Cell Signaling Technologies 14117S), Kme3 1:10,000 (Cell Signaling Technologies 14680S), AHCY 1:10,000 (ProteinTech 10757-2-AP). After washing in PBS with 0.01% Tween20, membranes were incubated with HRP-conjugated secondary antibodies for 1 h at room temperature. Protein bands were visualized using ECL substrate and imaged on a chemiluminescence detection system. See **Supplementary Table S3** for detailed list of all primary antibodies used in this study/

### Histone PTM analysis

Histones were acid-extracted from cultured cells as described previously^76^. Briefly, cells were collected at approximately 5 × 10^6^ cells/mL, pelleted by centrifugation at 300 × g for 10 min, washed with 1X PBS, and pelleted again at 300 × g for 10 min. Cell pellets were resuspended in hypotonic lysis buffer containing 10 mM Tris-HCl (pH 8.0), 1 mM KCl, 1.5 mM MgCl_2_, and 1 mM DTT, supplemented immediately before use with 1× protease and phosphatase inhibitors, and incubated for 30 min at 4°C with rotation. Nuclei were pelleted at 3,000 × g for 10 min at 4°C, washed once with hypotonic lysis buffer, and resuspended thoroughly in 0.4 N sulfuric acid. Samples were rotated for at least 30 min at 4°C to extract histones, then centrifuged at 16,000 × g for 10 min at 4°C to remove insoluble debris. The supernatant was transferred to a fresh tube, and histones were precipitated by dropwise addition of 71% (w/v) trichloroacetic acid to a final concentration of 33%, followed by incubation on ice for 30 min. Precipitated histones were collected by centrifugation at 16,000 × g for 10 min at 4°C, washed twice with 200 uL ice-cold acetone, with centrifugation at 16,000 × g for 10 min at 4°C after each wash, and then air-dried at room temperature for 20 min. Before derivatization, histones were resuspended in 50 mM ammonium bicarbonate (pH 8) containing 20% acetonitrile. In the fume hood, 2 µL of propionic anhydride and 10 µL of ammonium hydroxide (all Sigma Aldrich) were added to the samples. The mixture was incubated for 5 min and the procedure was repeated. Histones were then digested with 1 µg of sequencing grade trypsin (Promega) (1:20, enzyme:sample) diluted in 50 mM ammonium bicarbonate, pH 8, and incubated overnight at 37ºC. The derivatization reaction was repeated to derivatize peptide N-termini. The samples were dried in the Speedvac and stored in - 80ºC. For desalting, samples were resuspended in 100 µL of 0.1% TFA and loaded into a 96-well filter plate (Orochem) packed with Oasis HLB C-18 resin (Waters), which was equilibrated using 100 µL of the same buffer. Samples were washed with 100 µL of 0.1% TFA, eluted with 70 µL of a buffer containing 60% acetonitrile and 0.1% TFA, and dried using SpeedVac. After desalting, samples were resuspended in 10 µL of 0.1% TFA and loaded onto a Dionex RSLC Ultimate 300, coupled online with an Orbitrap Fusion Lumos (all Thermo Scientific). Chromatographic separation was performed using a two-column system, consisting of a C-18 trap cartridge (300 µm ID, 5 mm length) and an analytical column (75 µm ID, 25 cm length) packed in-house with reversed-phase Repro-Sil Pur C18-AQ 3 µm resin. Histone peptides were separated using a 60 min gradient from 4–30% buffer B (buffer A: 0.1% formic acid, buffer B: 80% acetonitrile + 0.1% formic acid) at a flow rate of 300 nL/min. The mass spectrometer was set to acquire spectra in a data-independent acquisition (DIA) mode. The full MS scan was set to 300–1100 m/z in the orbitrap with a resolution of 120,000 (at 200 m/z) and an AGC target of 5 × 105. MS/MS was performed in the orbitrap with sequential isolation windows of 50 m/z with an AGC target of 2 × 105 and an HCD collision energy of 30.

Raw files were imported into EpiProfile 2.0 software^77^. From the extracted ion chromatogram, the area under the curve was obtained and used to estimate the abundance of each peptide. To achieve the relative abundance of PTMs, the sum of all different modified forms of a histone peptide was considered as 100% and the area of the particular peptide was divided by the total area for that histone peptide in all of its modified forms. The relative ratio of two isobaric forms was estimated by averaging the ratio for each fragment ion with different mass between the two species. The resulting peptide lists generated by EpiProfile were exported to Microsoft Excel and further processed for a detailed analysis (**Supplementary Table S4**).

### RNA-seq

After 5 days of growth, HepG2 and MIHA cells were washed with PBS, harvested with Zymo DNA/RNA Shield reagent, and shipped at room temperature to Plasmidsaurus for library preparation and sequencing. Bulk 3′ RNA sequencing was performed using the Plasmidsaurus RNA-seq service. Libraries were generated using a 3′-end counting workflow in which poly(A)+ RNA was captured with an oligo-dT primer containing a sample barcode and unique molecular identifier, followed by reverse transcription, second-strand synthesis, tagmentation, and Illumina library amplification with unique dual indexes. Sequencing was carried out on an Illumina platform, and transcript abundance was quantified from deduplicated 3′ counting reads. Because the assay uses UMI-based 3′ end counting, it is optimized for transcriptome-wide gene-level differential expression analysis rather than transcript isoform discrimination. Plasmidsaurus returned raw FASTQ files, aligned deduplicated BAM files, QC summaries, and gene-count matrices for downstream analysis. Gene count matrices were analyzed for differential transcript abundance using the DESeq2 Bioconductor R package^78^ (**Supplementary Table S5**). Pathway analysis was performed with the ClusterProfiler Bioconductor R package^79^.

### Statistical analysis

All immunoblots were performed in at least two independent biological replicates. Metabolomics, RNA-seq, and all other relevant assay were performed with at least three independent biological replicates. Statistical analyses were performed either using GraphPad Prism (v10.1.0) or R (v4.4.0). Independent t-tests were performed to compare means between only two groups.

All plasmid constructs used in this study are detailed in **Supplemental Table S6**.

## Supporting information

Supplemental Figures S1 through S6

## Acknowledgments

This work was supported by the Hevolution Foundation Grant HF-GRO-23-1199128-32 to D.S., E.G., and D.M.H. This work was also supported by the National Institutes of Health [R01GM108646 to D.S., T32GM149364 to I.K].

Crystallographic data collection was performed at the Advanced Photon Source, a U.S. Department of Energy (DOE) Office of Science user facility operated for the DOE Office of Science by Argonne National Laboratory under Contract No. DE-AC02-06CH11357. Use of the Lilly Research Laboratories Collaborative Access Team (LRL-CAT) beamline at Sector 31 of the Advanced Photon Source was provided by Eli Lilly and Company, which operates the facility. The Einstein Crystallographic Core X-ray diffraction facility is supported by NIH Shared Instrumentation Grant S10 OD020068 and Cancer Center Support Grant P30 CA013330, which we gratefully acknowledge.

The mass spectrometry proteomics data have been deposited to the ProteomeXchange Consortium via the PRIDE^80^ partner repository with the dataset identifier PXD078525.

Some figures were created with BioRender. We are grateful to Rajesh K. Harijan for initial assistance with crystallography.

RNA-seq data is pending accession number to the Gene Expression Omnibus^81^

## Author contributions

I.K., D.S., D.M.H. conceived of the study. I.K., S.H., D.S., M.B., R.E., D.G. conceived of experiments. I.K., O.W.M, S.H., M.K.B., H.H. B.M.L. performed experiments. I.K, D.S., O.W.M, E.G., J.B., B.M.L. performed data analysis. J.A., S.G., & S.S. assisted with proteomics experiments. I.K. and D.S. wrote the manuscript. All authors approved the manuscript.

## AI Declaration

To improve concision in portions of this manuscript, during the preparation of this work the authors used ChatGPT. After using this tool, the authors fully reviewed and edited the content as needed and take full responsibility for the content of the publication.

