## Supplemental Figures S1 through S6 for "The GNMT N-terminus Couples Folate Feedback to Methyl-donor Homeostasis"

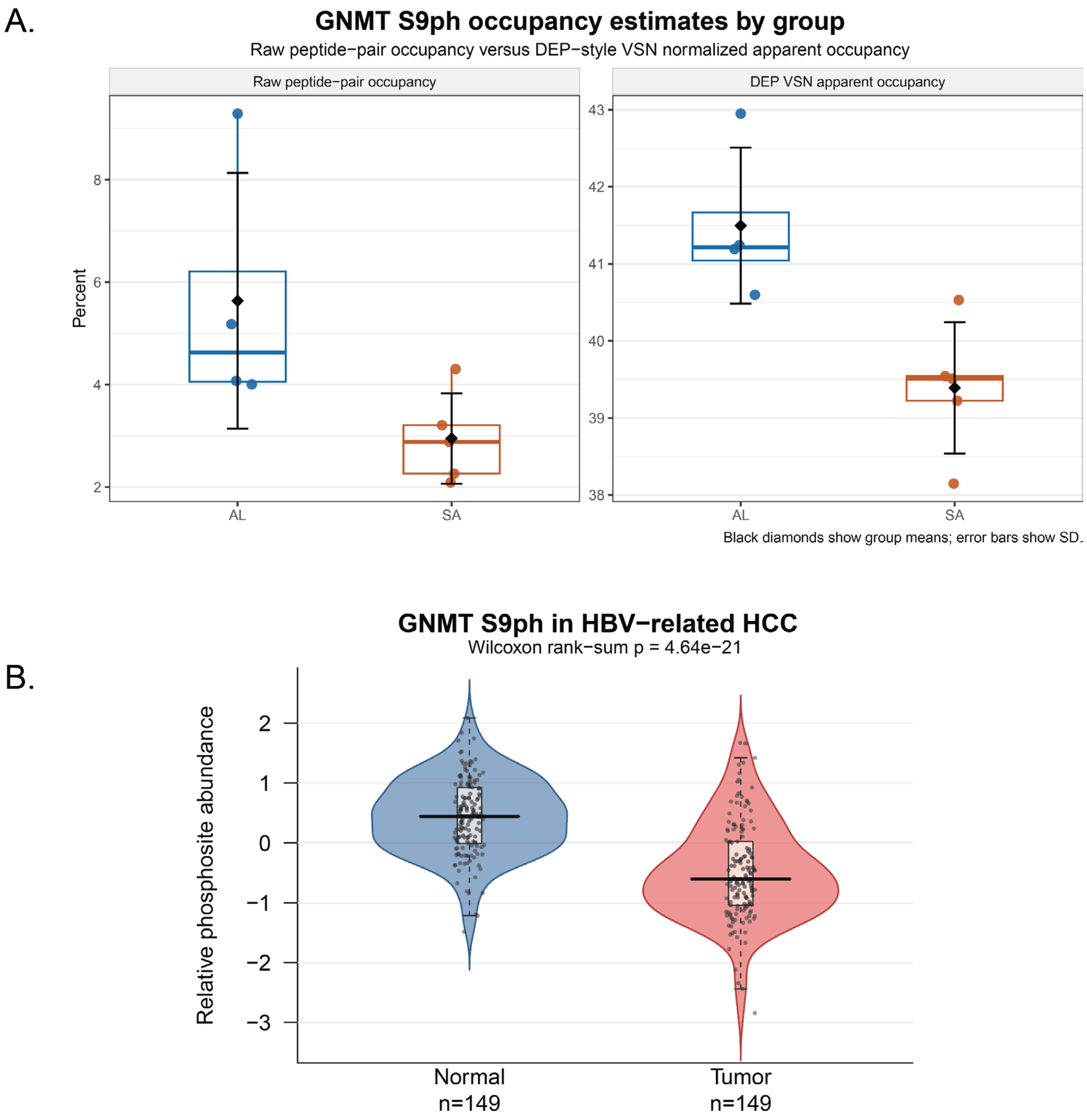


**Supplementary Figure S1. GNMT S9 phosphorylation (S9ph) varies across dietary and disease contexts. (A)** GNMT S9ph occupancy estimates in mouse liver samples from ad libitum-fed (AL) and sulfur amino acid-restricted (SA) mice. The left panel shows raw peptide-pair occupancy estimates, and the right panel shows DEP-style VSN-normalized apparent occupancy estimates. Individual points represent biological samples, black diamonds indicate group means, and error bars indicate standard deviation. **(B)** Relative GNMT S9ph abundance in matched normal and tumor samples from an HBV-related hepatocellular carcinoma proteogenomic dataset. Violin plots show the distribution of relative phosphosite abundance across normal and tumor groups, with overlaid boxplots and individual sample points; the Wilcoxon rank-sum test P value is shown above the plot.

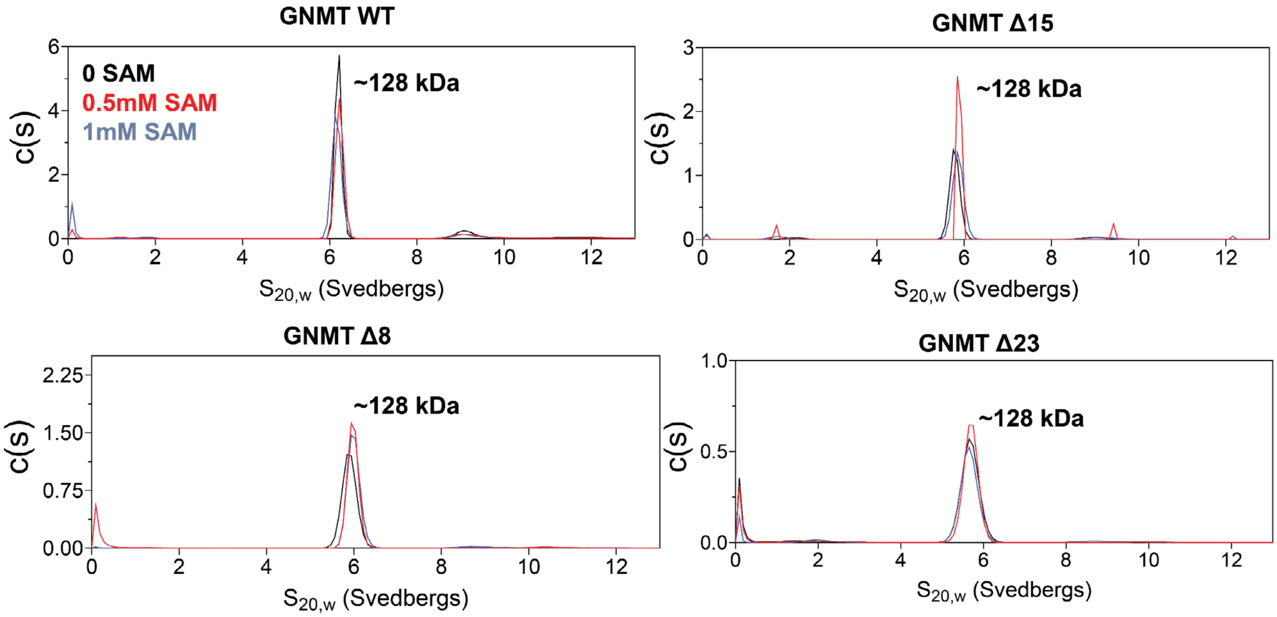


**Supplementary Figure S2. Analytical ultracentrifugation analysis of GNMT N-terminal truncation variants.** Sedimentation velocity analytical ultracentrifugation profiles of recombinant GNMT full-length, Δ8, Δ15, and Δ23 proteins measured in the presence of 0, 0.5, or 1 mM SAM. Each panel shows the sedimentation coefficient distribution for the indicated GNMT construct, with traces corresponding to the two SAM concentrations. This figure corresponds to the analytical ultracentrifugation experiments described in the "The N-terminus supports activity and SAM stability" section, which assess the oligomeric state of full-length and N-terminally truncated GNMT variants under SAM-bound conditions.

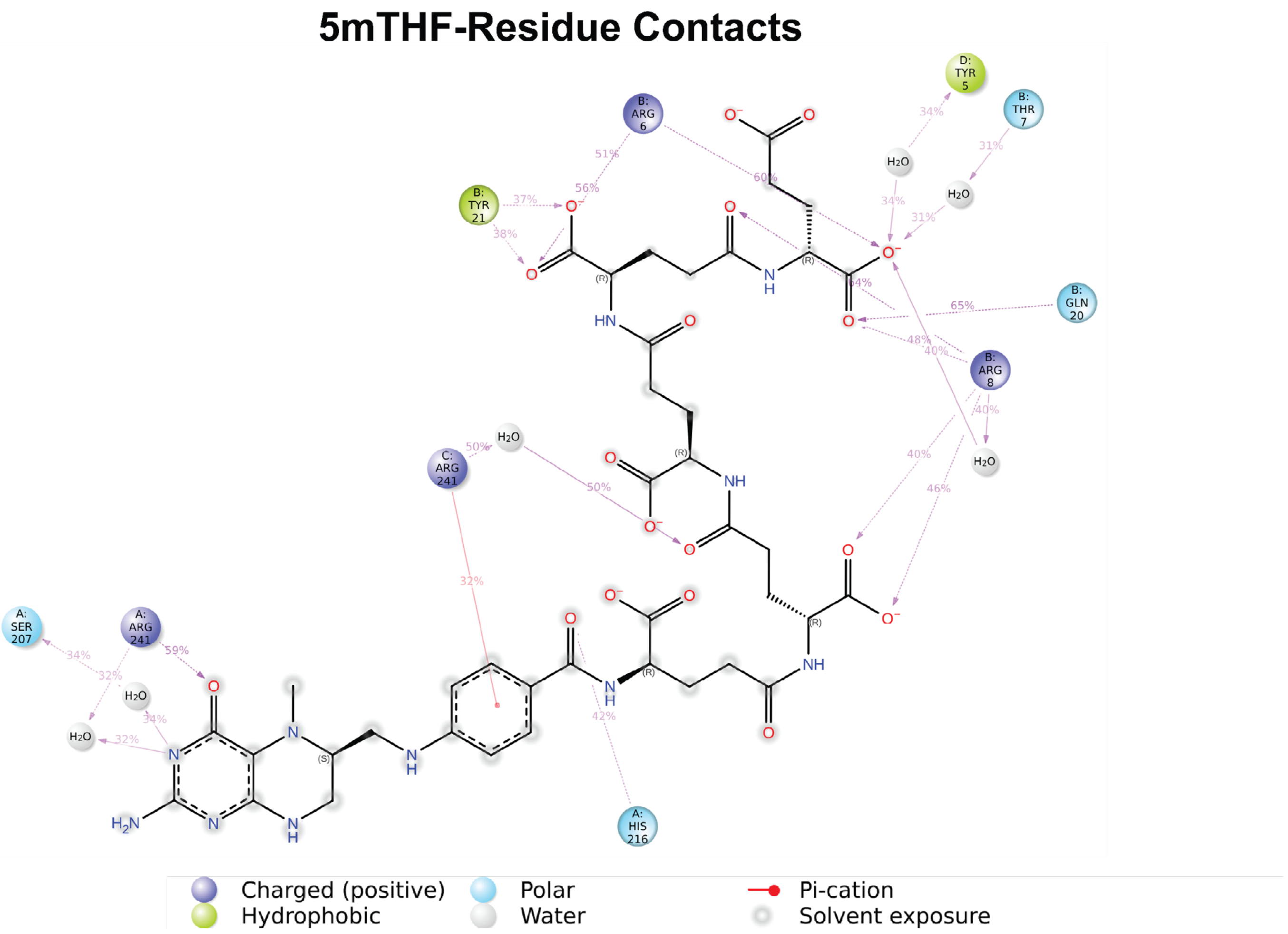


**Supplementary Figure S3. Residue contacts with 5mTHF in the GNMT-binding pocket.** Ligand interaction diagram showing atomic contacts between 5mTHF-(Glu)$_5$ and GNMT residues during the selected molecular dynamics trajectory. Interactions shown occurred for more than 30.0% of the simulation time over the selected trajectory from 0 to 1000 ns. Interacting residues and waters are labeled by chain and residue number, and dashed interaction lines indicate the interaction type and contact frequency. Interaction frequencies may exceed 100% when a single residue forms multiple simultaneous interactions with the same ligand atom. For example, arginine side chains can have multiple hydrogen-bond donors that contact a single hydrogen-bond acceptor. Residue classes and interaction types are indicated in the legend, including charged, hydrophobic, polar, water-mediated, pi-cation, and solvent-exposure interactions. This figure corresponds to the molecular dynamics analysis of the GNMT–5mTHF binding pocket described in the “S9ph is poised to disrupt the 5mTHF binding pocket” section, including contacts involving R241 and neighboring N-terminal residues.

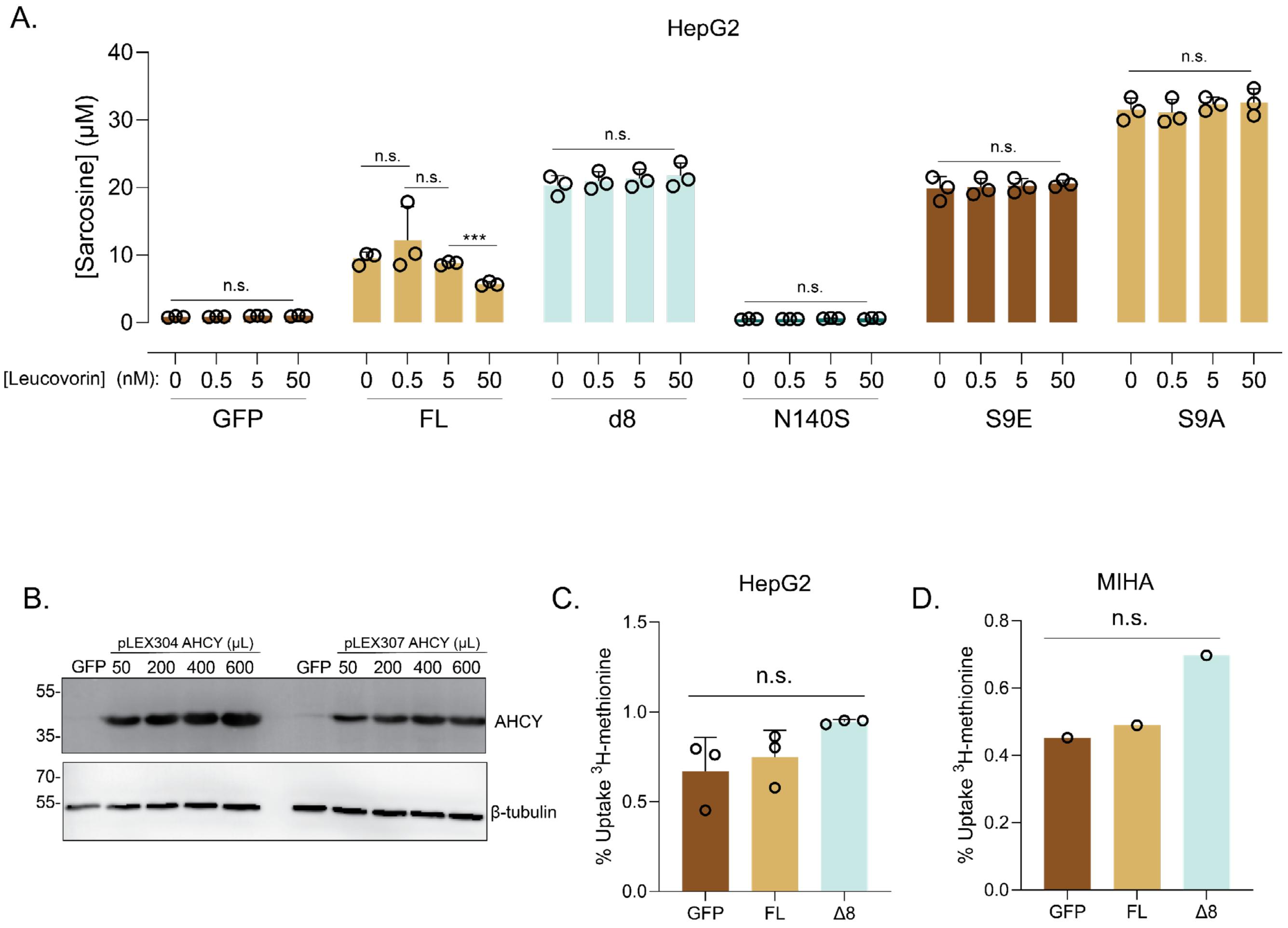


**Supplementary Figure S4. Leucovorin dose response, methionine uptake, and metabolite measurements in GNMT-expressing cells. (A)** Full leucovorin dose-response analysis of media sarcosine concentration in HepG2 cells expressing GFP control, full-length GNMT, GNMT Δ8, or GNMT S9E. Individual points indicate replicate measurements. **(B)** Western blot showing titration of pLEX304/07 AHCY lentivirus. The pLEX304 200uL condition was chosen for relevant experiments. **(C)** Cellular methionine uptake assay in HepG2 cells expressing GFP, full-length GNMT, or GNMT Δ8 after incubation with radiolabeled $^3$H-methionine. **(D)** Cellular methionine uptake assay in MIHA cells expressing GFP, full-length GNMT, or GNMT Δ8 after incubation with radiolabeled $^3$H-methionine.

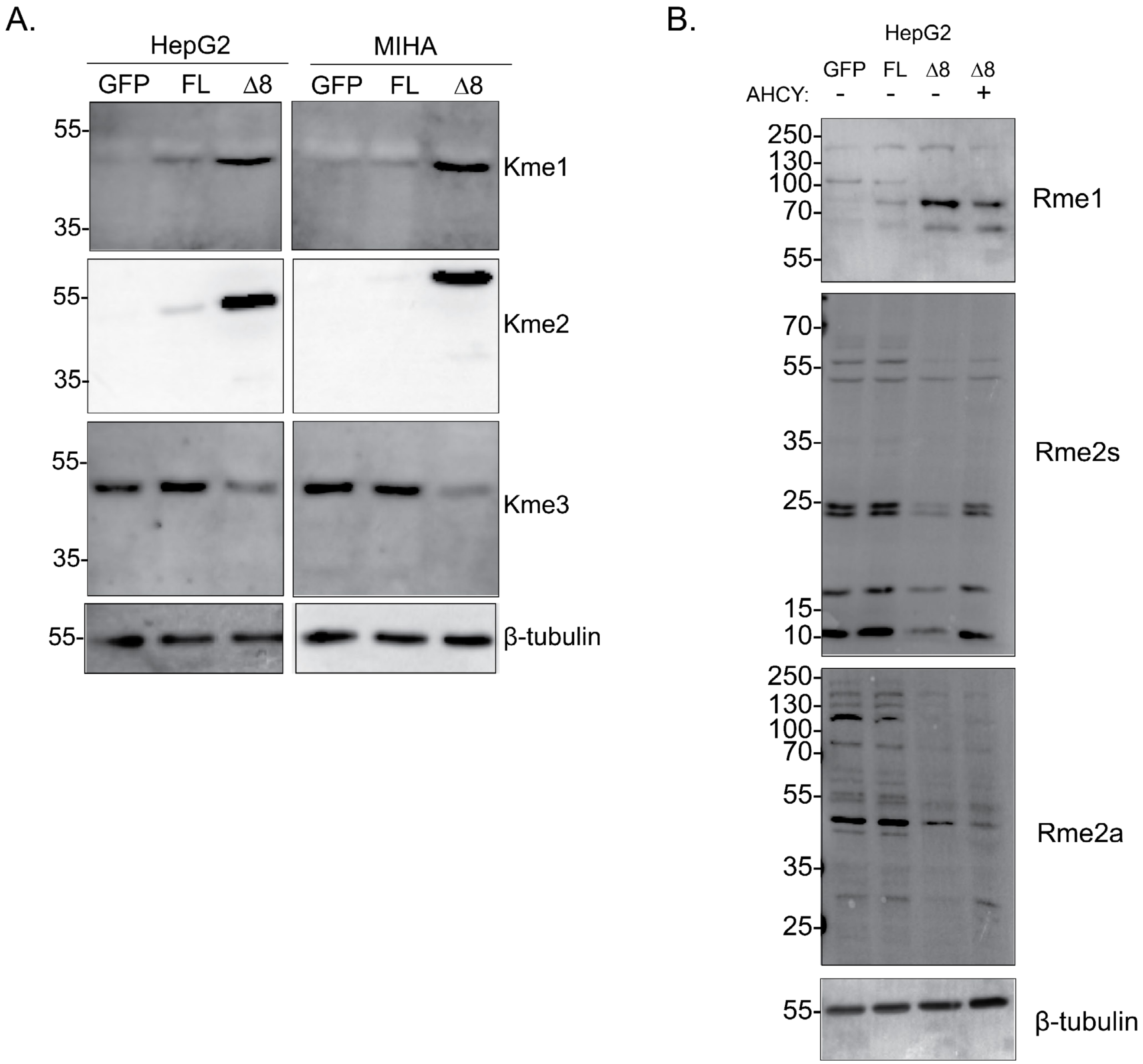


**Supplementary Figure 5. GNMT Δ8 expression alters global lysine methylation patterns.** Immunoblot analysis of global lysine methylation marks in HepG2 and MIHA cells expressing GFP control, full-length GNMT, or GNMT Δ8. Blots show pan-monomethyl lysine (Kme1), dimethyl lysine (Kme2), and trimethyl lysine (Kme3) signal across expression conditions, with β-tubulin loading controls.

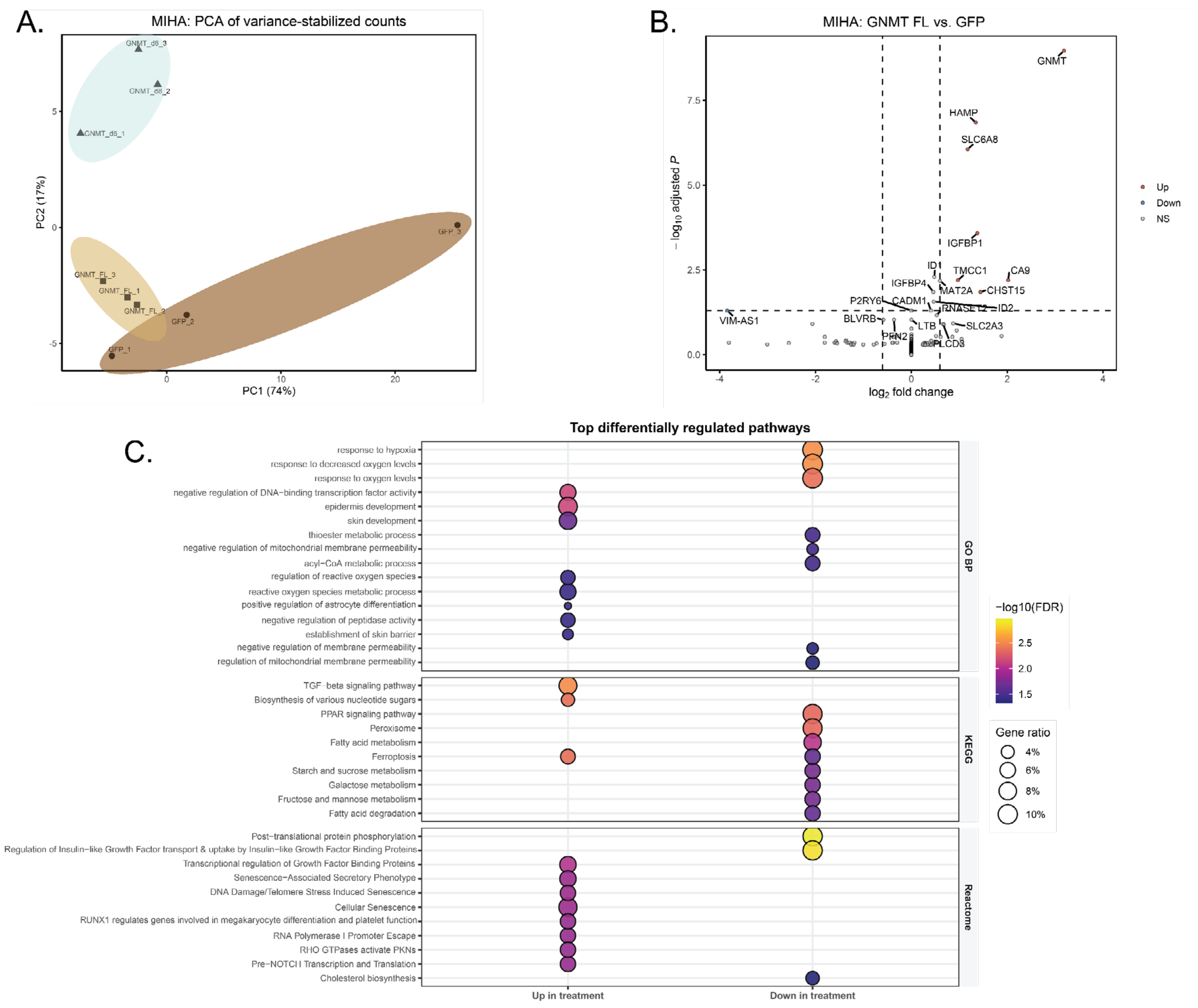


**Supplementary Figure 6. Transcriptomic remodeling induced by GNMT Δ8 expression in MIHA cells. (A)** Principal component analysis of variance-stabilized RNA-seq counts from MIHA cells expressing GFP control, full-length GNMT, or GNMT Δ8. Each point represents an individual biological replicate, and shaded ellipses indicate the distribution of replicates within each expression condition. **(B)** Volcano plot of differential gene expression in MIHA cells comparing full-length GNMT with GFP control, with log2 fold change on the x-axis and −log10 adjusted *P* value on the y-axis. Upregulated, downregulated, and non-significant genes are indicated, with selected genes labeled. **(C)** Dot plot of pathway enrichment analysis from MIHA GNMT Δ8 versus full-length GNMT comparison, grouped by GO Biological Process, KEGG, and Reactome categories. Dot size indicates gene ratio, and color indicates −log10 FDR.
